# TSC2 is a stress granule suppressor

**DOI:** 10.64898/2026.05.21.726761

**Authors:** Yizhe Ma, Natalie G. Farny

**Affiliations:** Department of Biology and Biotechnology, Worcester Polytechnic Institute, Worcester, MA, USA; Program in Bioinformatics and Computational Biology, Worcester Polytechnic Institute, Worcester, MA, USA

**Keywords:** Stress granules, RNAi screen, tuberous sclerosis complex, translational control

## Abstract

Stress granules (SGs) are dynamic, membrane-less ribonucleoprotein assemblies that form through liquid-liquid phase separation to prioritize stress-survival proteostasis. Through reanalysis of a *Drosophila* genome-wide RNAi screen, we identified a set of conserved SG suppressor genes and validated the top candidate, Tsc2, in both mouse and human cell lines. We illustrate that the complete loss of *Tsc2* leads to spontaneous, canonical, and translation-dependent SGs driven by mTORC1 hyperactivation in mouse embryonic fibroblasts (MEFs). In addition, the *Tsc2*-deficient MEFs also sensitized to endoplasmic reticulum stress, delaying SG clearance. In human cell lines, the siRNA-mediated partial reduction of *TSC2* in U2OS cells, and in human tuberous sclerosis patient fibroblasts, does not induce spontaneous SGs. Instead, the sensitivity to ER stress, translation perturbation, and delay in clearance correlate with the remaining levels of TSC2, suggesting that TSC2 functions as a threshold-dependent regulator of SG assembly. Together, our findings provide a comprehensive list of novel conserved SG regulators and establish TSC2 as a key regulator of SG dynamics.

## INTRODUCTION

Stress granules (SGs) are cytoplasmic, dynamic, and membrane-less aggregates of proteins and RNAs that form through liquid-liquid phase separation (LLPS) in response to a variety of environmental and biotic stressors (Parobkova and Matej, 2021, Li et al., 2024, Kamagata et al., 2022). While classically viewed as passive storage depots that sequester translation factors to inhibit protein synthesis, recent evidence suggests SGs enhance the integrated stress response by prioritizing the translation of stress-survival transcripts and protecting small ribosomal subunits (40S) and pre-initiation complexes from degradation during stress (Grao-Cruces et al., 2023, Smith and Bartel, 2026, Kedersha et al., 2000, Wu et al., 2025). SG assembly is a tightly regulated and reversible process; however, failure in SG clearance upon stress relief can result in the formation of cytotoxic, pathogenic aggregates (Polymenidou and Cleveland, 2011, Baron et al., 2013, Park et al., 2025, Huang et al., 2025, Millar et al., 2023).

Efficient disassembly of SGs after stress removal is critical for maintaining cellular proteostasis. SG clearance is facilitated by chaperones, autophagy, and AAA+ ATPase Valosin-containing protein (VCP), which extracts ubiquitinated proteins to prevent accumulation misfolded protein. (Hofmann et al., 2019, Ramaswami et al., 2013, Gan et al., 2018, Hu et al., 2022). Inhibiting VCP function impairs SG disassembly, results in the persistence granules after stress has been removed. Persistent SGs or defects in SG disassembly are implicated in neurodegenerative diseases including Amyotrophic Lateral Sclerosis (ALS) and Frontotemporal Dementia (FTD) (Ma and Farny, 2023, Ishaq and Russell, 2025, Ueda et al., 2024, Li et al., 2022, Nedelsky and Taylor, 2022, Cui et al., 2024). Genetic depletion of key factors involved in SG clearance result in the formation of spontaneous SGs (Marcelo et al., 2021, Yuan et al., 2025, Nahm et al., 2020, Becker et al., 2017, Bakthavachalu et al., 2018, Maharjan et al., 2017, Zheng et al., 2022, Chitiprolu et al., 2018, Chew et al., 2019, Seguin et al., 2014).

The TAR DNA binding protein 43 (TDP-43) is an RNA binding protein mainly present in the nucleus that regulates RNA metabolism of mRNAs encoding proteins that play roles in neurodegeneration (Dewey et al., 2012). TDP-43 is a key regulator of SG dynamics following stress by stabilizing SG core components including G3BP1 and TIA-1. As a hallmark of ALS/FTD, the nuclear depletion of TDP-43 or specific pathological mutations impair SG assembly (McDonald et al., 2011). The role of SGs in TDP-43 transition from dynamic liquid droplets to insoluble inclusions is under debate. The seeding hypothesis suggests that SGs act as seeds, where chronic stress causes SGs which serve as a hub that locally concentrates TDP-43 to trigger a liquid-to-solid phase transition (Dewey et al., 2012, Liu-Yesucevitz et al., 2010, Rummens et al., 2025, Scialò et al., 2025). Alternatively, others hypothesized that TDP-43 aggregation is independent of SG formation since the high concentration of TDP-43 is sufficient to exceed the threshold for LLPS (Dewey et al., 2012, Streit et al., 2022, Hans et al., 2020, Fernandes et al., 2020). Recent evidence suggests that SG formation is an intermediate step of TDP-43 aggregation (Yan et al., 2025, De Boer et al., 2021). In this proposed mechanism, SGs initially nucleate TDP-43 recruitment, and then TDP-43 transitions to a dense, solid subphase inside SGs. As TDP-43 aggregates mature and stabilize, the initial SG scaffold dissociates or degrades, ultimately dissociating SGs from TDP-43 aggregates. The hypothesis requires additional evidence, and the potential role of any other genetic factors remains unknown.

Multiple studies have identified regulators of SGs and related ribonucleoprotein (RNP) granules. Early work investigating RNA triage (Anderson and Kedersha, 2008) and genetic screens in lower eukaryotes defined conserved regulatory networks, including granulophagy for SG clearance in *S. cerevisiae* (Buchan et al., 2013) and P-body assembly pathways in *C. elegans* (Sun et al., 2011). In mammalian systems, studies have largely been targeted to specific gene subsets. RNAi screening of the druggable genome linked the hexosamine biosynthetic pathway and O-GlcNAc modification to SG assembly under arsenite treatment (Ohn et al., 2008). Systems-level screening approaches have identified regulators of membrane-less organelle (Berchtold et al., 2018), specific RNA-binding proteins (Wheeler et al., 2020), and cell-cycle kinases (Haneke et al., 2020). Yang et al., proposes that SGs are regulated by a multivalent protein-RNA network and identifies extrinsic factors that either strengthen this assembly through positive cooperativity or weaken it through competitive inhibition (Yang et al., 2020). Other studies targeted germ cell specific factors DAZL (Kim et al., 2012), and translation initiation factors (Mokas et al., 2009), mapped the SG proteome to identify SG regulators required for assembly (Youn et al., 2018), or defined the molecular mechanisms of assembly and disassembly (Hofmann et al., 2021).

Despite the fact that these prior studies have defined key SG regulators, a genome-wide identification of conserved genetic suppressors of SG assembly remains lacking. To address this gap, we performed an analysis of a high-throughput whole genome RNAi screen in *Drosophila* S2R+ cells to identify novel SG suppressor genes. Our screen identified 271 *Drosophila* candidate poly(A)^+^ RNAs granule suppressor genes, corresponding to 365 human orthologs. Among the top hits, the tuberous sclerosis complex 2 gene (TSC2) was identified as a conserved regulator of SG assembly. The TSC complex consists of TSC1, TSC2, and TBC1D7, and functions as a key negative regulator of mechanistic target of rapamycin complex 1 (mTORC1) by inhibiting Rheb-dependent kinase activation (Dibble et al., 2012, Santiago Lima et al., 2014, Qin et al., 2016, Mallela and Kumar, 2021, Karalis et al., 2024). The relationship of mTORC1 and SGs is a negative feedback loop; stress-induced mTORC1 inhibition facilitates SG formation by allowing ribosomes to dissociate from mRNA (Cadena Sandoval et al., 2021). We propose that dysregulation of the TSC-mTORC1 axis leads to a proteostatic imbalance, causing impaired assembly and clearance (McCubrey et al., 2012, Xing et al., 2023).

Using genetic and pharmacological methods in mouse and human cell lines, we illustrate that only the complete loss of *TSC2* leads to spontaneous SGs. The sensitivity of *TSC2*-depleted cells to endoplasmic reticulum (ER) stressors and translation manipulation correlates to the level of TSC2 remaining in the cell. Our findings establish TSC2 as a key regulator of SG dynamics and reveal a pathway-specific sensitivity linking mTORC1 hyperactivation to proteostatic imbalance. Together, our findings establish TSC2 as a key regulator of SG dynamics and provide a valuable resource for further dissection of pathways controlling SG assembly and clearance.

## MATERIALS AND METHODS

### Re-analysis of a genome-wide RNA interference (RNAi) screen to identify potential SG regulators

A previously published genome-wide RNAi performed in *Drosophila* S2R+ cells targeting 13,620 genes was used as the source dataset for this study (Farny et al., 2008) (Figure 1A). In the original screen, cells were treated with double-stranded RNAs (dsRNAs) and subjected to fluorescence in situ hybridization (FISH) using an Oligo-dT(30) probe to visualize poly(A)^+^ RNA distribution, as previously described (Farny et al., 2008).

**Figure 1.**
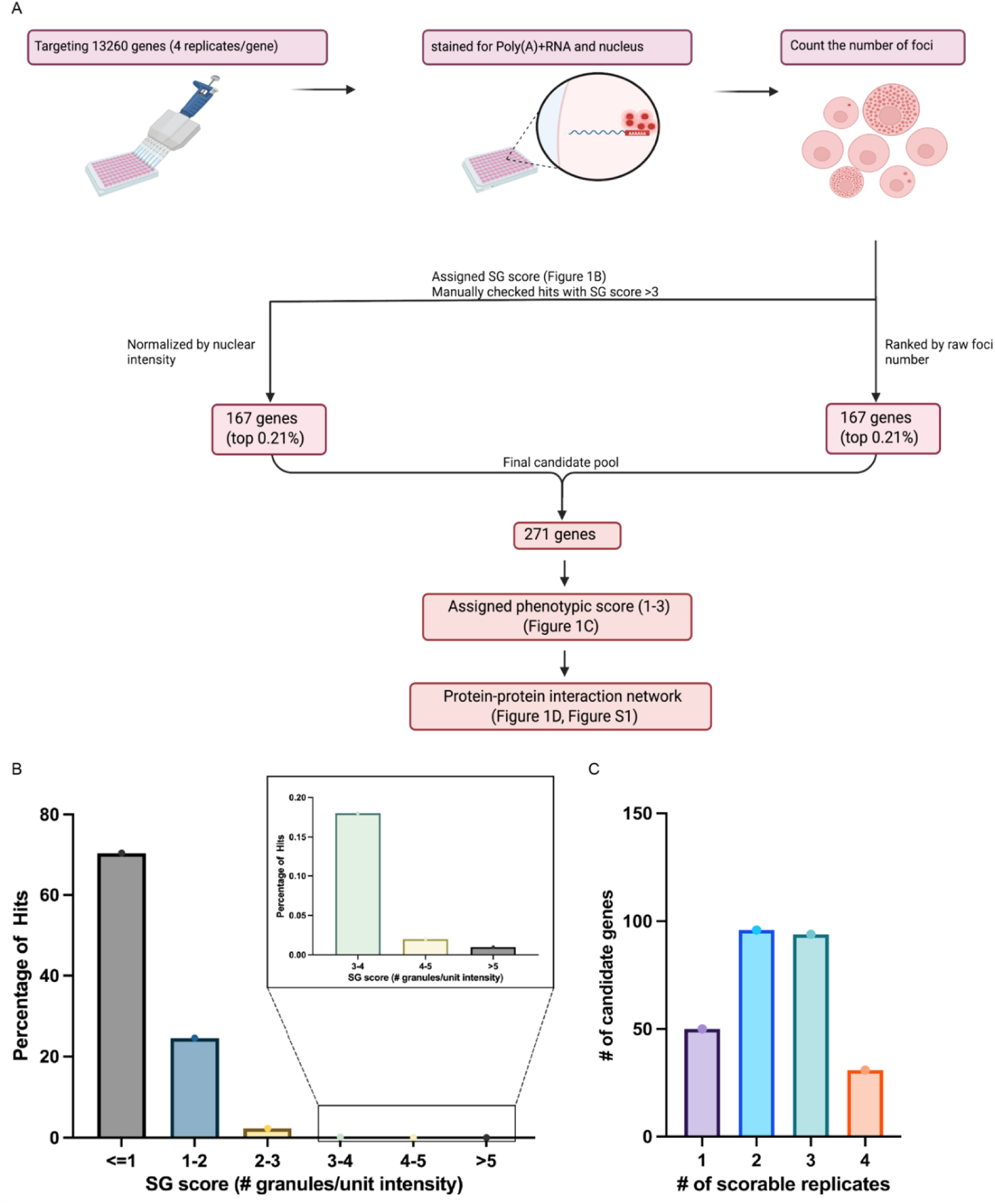
A whole genome *Drosophila* RNAi screen for suppressors of poly(A)^+^ RNA granules. A) A whole genome RNAi screening was conducted in *Drosophila* S2R+ cells stained with Cy3-labeled oligo-dT(30) for total poly(A)^+^ RNA granules. The number of poly(A)^+^ RNA granules were counted by CellProfiler and then normalized to the intensity of the corresponding DAPI channel. B) SG score distribution. C) Distribution of number of genes with number of scorable replicates.

To identify potential stress granule (SG) suppressor genes, we re-analyzed the imaging dataset by quantifying poly(A)^+^ RNA foci using CellProfiler. For each gene, an SG score was calculated by normalizing the number of cytoplasmic poly(A)^+^ RNA foci to the intensity of the nuclear channel (Figure 1B). Candidate genes with SG scores greater than 3 granules per unit intensity (top 0.21%) were manually inspected, and 167 genes were selected for phenotypic classification (Table S1).

To ensure that strong candidates were not excluded due to the normalization procedure, we also select the top 0.21% candidates based on the raw number of poly(A)^+^ RNA foci per image without normalization (Table S1). Combining candidates identified by both approaches yielded 271 genes that were manually assigned phenotypic scores ranging from 1 to 3 based on the severity of the SG phenotype (Figure 1C; Table S1).

### Protein-Protein interaction network

Protein-Protein interaction (PPI) network analysis for 271 *Drosophila* genes was performed using the STRING database (Figure S1) (Szklarczyk et al., 2023). Gene ontology (GO) term analysis was conducted using STRING database to identify biological processes associated with the identified genes (Szklarczyk et al., 2023).

PPI network analysis for human orthologs of 271 Drosophila genes was performed using Human Reference Interactome (HuRi) (Figure 2) (Luck et al., 2020). The network was visualized using Cytoscape (v3.10.4) (Shannon et al., 2003). Nodes were organized using functional clustering.

**Figure 2.**
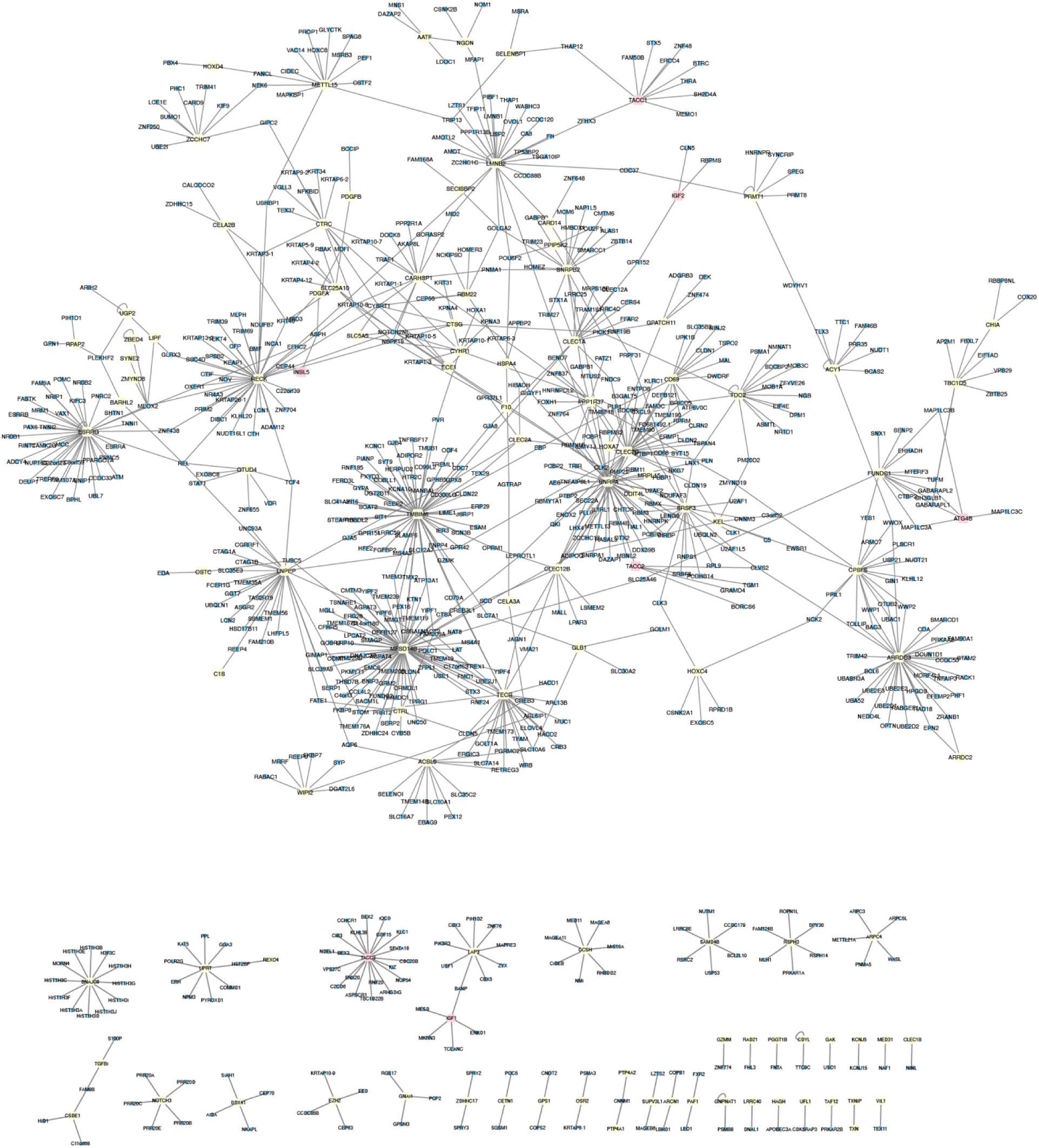
Comparative interactome analysis of identified stress granule suppressors. The human orthologs of the 271 *Drosophila* genes were mapped using the Human Reference Interactome (HuRi) database. Yellow nodes represent the human orthologs of the original *Drosophila* screen hits. Pink nodes represent the human orthologs of the top *Drosophila* candidate. Blue nodes represent additional interacting proteins identified by the HuRi pipeline. All interactions in this panel are derived from a systematic experimental screening pipeline, possessing a data quality and fraction correct comparable to high-quality literature-curated binary interactions with at least two pieces of independent experimental evidence.

### Cell culture

*Tsc1^−/−^* and *Tsc2^−/−^* MEF cells and wild type controls were generously provided by Dr. David Kwiatkowski of Havard Medial School (Kwiatkowski et al., 2002, Zhang et al., 2003, Onda et al., 1999). U2OS cells (ATCC HTB-96) and MEF lines were maintained at 37°C in a CO_2_ incubator (5% CO_2_) in DMEM (Gibco^TM^, cat no: 11965-084) supplemented with 10% FBS (Gibco^TM^, cat no: A5256701), 1% sodium pyruvate (Gibco^TM^, cat no: 11360070), and 1% penicillin/ streptomycin (Gibco^TM^, cat no: 15140122). Patient fibroblast cells (AHI: GM07522 and TSP: GM04520) were obtained from Coriell Institute for Medical Research. AHI and TSP were maintained at 37°C in a CO_2_ incubator (5% CO_2_) in EMEM (ATCC^TM^, cat no: 8111924) supplemented with 15% FBS (Gibco^TM^, cat no: A5256701).

### Drug treatment

Cells were treated with Puromycin (Gibco^TM^, cat no: A11138-03), cycloheximide (Themo Scientific, cat no: J66004.XF), thapsigargin (Millipore Sigma T9033), MG132 (Millipore Sigma 474790), sodium arsenite (Sigma, S7400), and DBeQ (Millipore Sigma SML0031) for indicated times.

### Transient transfection

U2OS cells were plated onto coverslips at 0.7×10^5^ cells and transfected with siRNA using Lipofectamine^TM^ RNAiMAX (Thermo Scientific^TM^, cat no:13778075) for knockdown or siRNA cotransfected with TDP43 NLS YFP construct (Addgene, cat no: 84912) using Lipofectamine^TM^ 3000 Transfection Reagent (Thermo Scientific^TM^, cat no: L3000015) following manufacture protocol. Following transfections cells were treated as indicated and prepared for immunofluorescence microscopy or western blot. The following siRNA were used: si*Control* (Dharmacon^TM^, cat no:D-001810-01-05) and si*TSC2* (Dharmacon^TM^, cat no: L-003029-00-0005)

### Immunofluorescence

1.5×10^5^ cells were plated into 12-well plates seeded with glass cover slips. The following day chemical treatments were applied as indicated. Following treatment, cells were fixed with 4% paraformaldehyde for 10 minutes, permeabilized with methanol for 10 minutes, and blocked for at least 1 hour with 5% BSA (Research Products International, cat no: 9048-46-8) in PBS. Primary antibody was diluted in blocking solution and incubated at room temperature for 1hour or at 4°C overnight. Coverslips were washed three times with 1X PBS and then incubated with secondary antibodies and Hoechst (Thermo scientific, cat no: 62249) for 1h at room temperature. Coverslips were washed 3 times and mounted on glass microscope slides using Vinol. The following primary antibodies were used: G3BP1 (Proteintec®, mouse cat no: 66486-1-Ig; Proteintec®, rabbit cat no: 13057-2-AP), eIF3η (Santa Cruz Biotechnology, cat no: sc-137214, eIF4E (Proteintec®, cat no: 11149-1-AP), and eIF4G (Proteintec®, cat no: 15704-1-AP). The following secondary antibodies from Jackson ImmunoReseach Laboratories INC. were used: Alexa Fluor ® 488-conjugated affiniPure Donkey Anti-Rabbit IgG (711-545-152), Alexa Fluor®594-conjugated affiniPure Donkey Anti-Rabbit IgG (711-585-152), Alexa Fluor®488-conjugated affiniPure Donkey Anti-Mouse IgG (715-545-150), and Alexa Fluor® 594-conjugated affiniPure Donkey Anti-mouse IgG (715-585-150).

### Microscopy and SG quantification

Fluorescence microscopy was performed using a Zeiss Axio Observer A1 inverted Fluorescence Microscope equipped with X-cite 120LED Boost High-Power LED illumination System. SG formation was quantified by manually scoring at least 250 cells and 3 fields of view under 40X magnification. Experiments were performed in technical duplicate with at least four independent biological replicates. The numbers of SG foci per cell were quantified by CellProfiler.

### Western blot

Protein extracts were prepared by adding 1X SDS sample buffer (62.5mM Tris (pH 6.8), 4% glycerol and 1.6% SDS) onto cells. Samples were boiled for 7 minutes at 95°C. DTT was added to each sample to make the final concentration to 16.7ng/ml and boiled for another 3 minutes. Samples were separated using tris-glycine polyacrylamide gels (Novex^TM^, cat no: XP04202BOX) in 1X tris-glycine running buffer (12.5mM tris base, 75mM glycine, and 1% SDS) Proteins were transferred onto nitrocellulose membranes (Thermo Scientific, cat no: 88018) in 1X transfer buffer (12.5mM tris base, 75mM glycine, and 20% methanol) at 110V. Membranes were blocked in 5% BSA in TBS-tween (TBS-T) for 2 hour at room temperature. Membranes were incubated with primary antibodies diluted 1:1000 TBS-T overnight at 4°C. Membranes were washed six times with TBS-T then incubated with HRP-conjugated secondary antibodies diluted at 1:5000 in TBS-T. Protein bands were detected using SuperSignal^TM^ West Pico PLUS Chemiluminescent Substrate (cat no: 1863095) and visualized on Azure c600 Imaging System. The following primary antibodies from Cell Signaling Technology were used: TSC1 (cat no: 6935T), TSC2 (cat no: 4308T), P-p70 S6 Kinase (cat no: 9208T), p70 S6 kinase (cat no: 2708T), and GAPDH (cat no: 97166S). The following secondary antibodies were used: Anti-rabbit IgG (cat no: 7074P2) and Anti-mouse IgG (cat no: 7076P2).

### Statistical analysis

Statistical analyses were performed using one-way or two-way ANOVA in GraphPad Prism (Version 10.6.1), as indicated in the figure legends. Statistical significance was defined as *P < 0.05, **P < 0.01, ***P < 0.001, and ****P < 0.0001. N indicates the number of independent biological replicate experiments.

## RESULTS

### Identification of stress granule suppressor genes

Poly(A)^+^ RNAs are defining components of many types of SGs, PBs, germ granules, neuronal granules, and other RNP granules (White and Lloyd, 2011, Kedersha et al., 1999, Anderson and Kedersha, 2009). To identify genes that suppress Poly(A)^+^ granule formation, we re-analyzed a previously published genome-wide RNAi imaging dataset generated in *Drosophila* S2R+ cells (Farny et al., 2008). We quantified Poly(A)^+^ RNA foci and assigned a SG score to each gene (Figure 1A and Figure 1B). This analysis identified 271 candidate genes (Table S1). To define functional relationship among these candidates, we performed protein–protein interaction (PPI) network analysis using the STRING database, based on both experimentally validated and predicted interactions (Figure S1A, Figure S1B, and Figure S2). This analysis resulted in an interconnected network composed of diverse functional clusters. To focus on direct associations, we next restricted the network, excluding all predicted interactions (Figure S1C and Figure S1D). Gene ontology (GO) analysis did not reveal any single dominant or significantly enriched pathway within the network, suggesting that SG regulation is not governed by a single pathway but rather emerges from multiple interconnected cellular processes (Figure S1B, Figure S1D, Figure S2B, and Figure S3). Nevertheless, several modest GO enrichments were observed, including groups associated with ribosome biogenesis and nucleolar function, basal transcription machinery, and stress response pathways, indicating that these functional modules represent localized contributions within a broader, distributed SG suppressor network.

To assess evolutionary conservation, *Drosophila* genes were mapped to their human orthologs and projected onto the Human Reference Interactome (HuRi), followed by visualization in Cytoscape (v3.10.4) (Figure 2) (Luck et al., 2020, Shannon et al., 2003). Despite the increased network complexity due to ortholog expansion, the resulting human interactome remained significantly connected, suggesting that SG suppressor pathways are conserved and maintain physical connectivity across species.

To refine our candidate list, we prioritized 67 candidate genes with the most robust and consistent phenotypes across all four replicate images (Table S2). The top-ranked genes, with the highest replication and phenotype scores, are *Ilp2*, *Atg4b*, *Tacc*, and *Gig* (Table 1). Among these hits, *Gig*, the *Drosophila* ortholog of *TSC2*, drew our attention. TSC2 functions with TSC1 and TBC1D7 in the conserved TSC complex, which negatively regulates mTORC1 signaling through inactivation of RHEB (Dibble et al., 2012, Santiago Lima et al., 2014, Qin et al., 2016, Mallela and Kumar, 2021). The TSC complex acts as a central node for AKT, WNT, ERK, AMPK, NFκB signaling pathway in response to growth factor stimulation, nutrient deprivation, hypoxia, and cytokines to modulate mTOR activity (Mallela and Kumar, 2021, Wang et al., 2025b, Huang and Manning, 2009, Zhang et al., 2003, Potter et al., 2002, Zhu et al., 2023, Wang et al., 2025a, Yuan et al., 2024). Mutation in *TSC1/2* leads to hyperactive mTOR which in turn drives the autosomal dominant disorder Tuberous Sclerosis Complex, a highly variable, multisystem disorder characterized by non-malignant, benign growths (hamartomas) in the brain, skin, kidneys, heart, and lungs (Mallela and Kumar, 2021, Wang et al., 2025b, Reyna-Fabián et al., 2020, Caban et al., 2016, Rosset et al., 2017). The previous literature has shown that *Tsc1 ^−/−^* and *Tsc2 ^−/−^* mouse embryonic fibroblast (MEF) cells exhibit increased basal mTORC1 activity, impaired autophagy, and enhanced cellular stress responses, including increased cellular apoptosis and phosphorylation of eIF2α upon ER stresses (Qin et al., 2010, Choi et al., 2018, Pal et al., 2019, Wu et al., 2010, Kang et al., 2011, Larsen et al., 2024, Alesi et al., 2024, Marqués et al., 2024, Patursky-Polischuk et al., 2009, Bjornsti and Houghton, 2004, Han et al., 2022, Danos et al., 2013). Notably, *Tsc2 ^−/−^* MEF cells reported to form increased numbers of SGs upon sodium arsenite and heat shock (Kosmas et al., 2021). Given these links between TSC2, mTOR signaling, and SGs, we further investigated *TSC2* as a compelling candidate for SG suppression.

**Table 1.** Top-ranked candidate potent SG suppressor genes. The top four candidate genes from the *Drosophila* RNAi screen were selected based on their consistent and robust phenotypic scores. The table lists the *Drosophila* gene symbol, full gene name, known biological function, and the corresponding human ortholog(s).

| Gene Symbol | Full gene name | Biological Function | Human Orthologs |
| --- | --- | --- | --- |
| *Ilp2* | Insulin-like peptide 2 | Maintains normoglycemia (a genetic strategy to measure circulating *Drosophila* insulin), determining adult lifespan; germ-line stem-cell niche homeostasis, and insulin receptor signaling pathway | *INSL5, IGF1, IGF2* |
| *Atg4b* | Autophagy-related 4b | Cysteine protease that regulates Atg8a activity by allowing Atg8a attach to membranes to form autophagosome and recycling Atg8a from membranes to regulate autophagic flux | *ATG4B, ATG4D* |
| *Tacc* | Transforming acidic coiled-coil protein | Centrosomal protein that stabilizes microtubules during cell division | *TACC1, TACC2, TACC3* |
| *Gig* | gigas | gigas (*gig*) encodes a tumor suppressor protein that, together with the product of *Tsc1*, controls cellular growth via antagonizing insulin and TOR signaling pathways | *TSC2* |

### Loss of *Tsc2* induces spontaneous, canonical, and mTOR-dependent SGs

To investigate the molecular role of *Tsc2* in regulating SG formation, we used both *Tsc1 ^−/−^* and *Tsc2 ^−/−^* mouse embryonic fibroblast (MEF) cell lines (Figure 3A). Consistent with the established role of the TSC complex as a negative regulator of mTORC1, both *Tsc1 ^−/−^*and *Tsc2 ^−/−^* cells exhibited upregulated phosphorylation of p70 ribosomal protein S6 kinase, suggesting hyperactive mTORC1 signaling (Figure 3A and Figure 3B).

**Figure 3.**
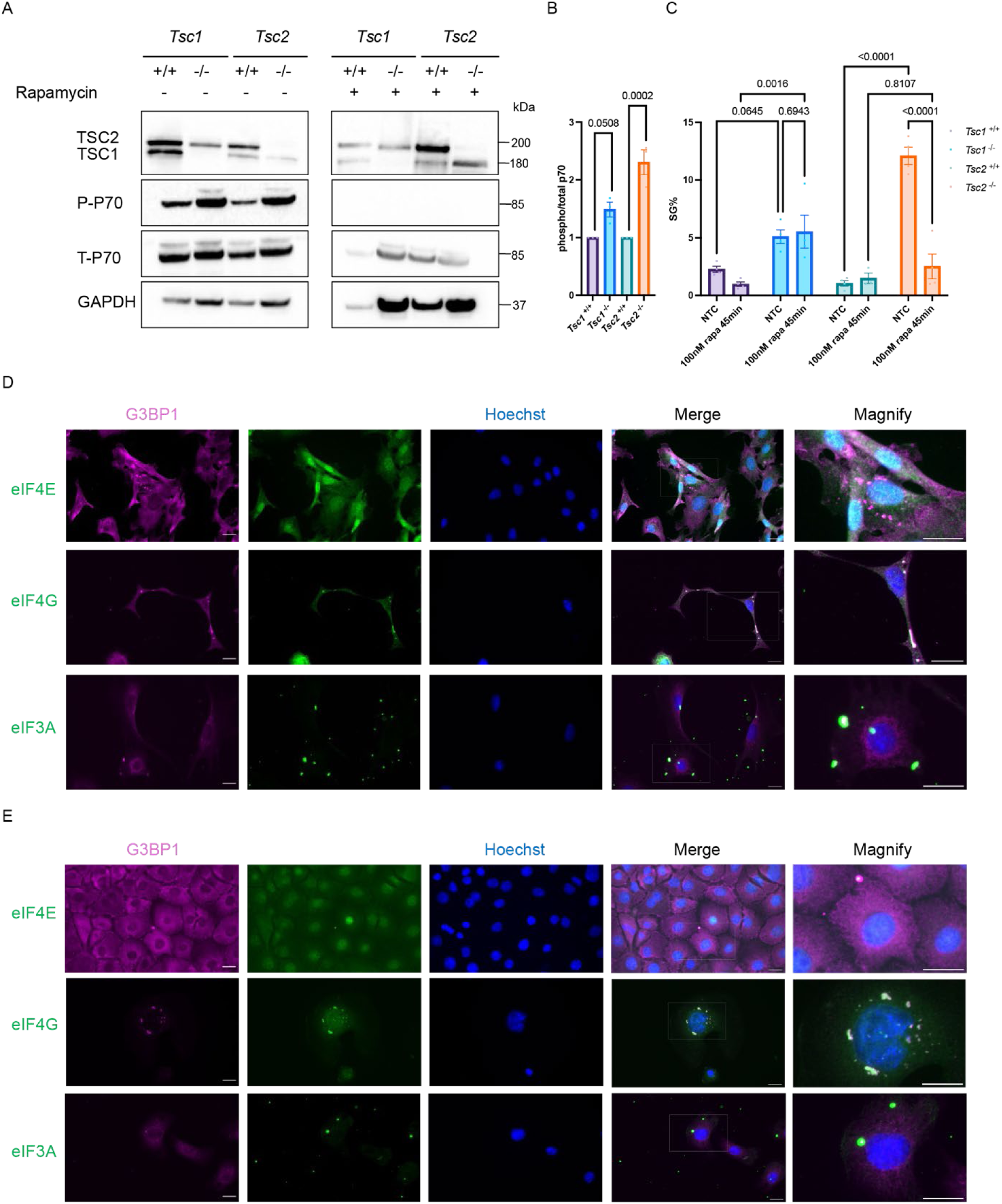
Characterization of *Tsc1 ^−/−^* and *Tsc2^−/−^* induced spontaneous SGs. A) Western blot analysis of Phospho-p70 S6 Kinase (P-P70) and Total p70 S6 Kinase (T-P70) in *Tsc1 ^−/−^* and *Tsc2 ^−/−^* MEF cells under untreated conditions or following rapamycin treatment. GAPDH was used as a loading control (n=3). B) Quantification of Phospho-p70 S6 Kinase (P-P70) and Total p70 S6 Kinase (T-P70) protein levels in untreated cells from the western blot shown in panel A (mean ± SEM, n=3). Statistical significance was determined using one-way ANOVA with Bonferroni’s multiple comparisons test. C) Quantification of the number of SGs formed in *Tsc1^−/−^* and *Tsc2 ^−/−^* MEF cells under rapamycin treatment (mean ± SEM, n=4). Statistical significance was determined using two-way ANOVA with Tukey’s multiple comparisons test. Colocalization analysis of SG marker G3BP1 and eukaryotic initiation factors for D) *Tsc1 ^−/−^* induced spontaneous SGs and E) *Tsc2 ^−/−^* induced spontaneous SGs. Scale bar, 20µM.

Based on our *Drosophila* RNAi screen results and phenotypic score ranking, we hypothesized that the genetic deletion of *Tsc2* would lead to spontaneous SG formation. Through immunofluorescence staining of the defining SG marker G3BP1, we confirmed that both *Tsc1^−/−^ and Tsc2 ^−/−^* cells formed spontaneous SGs, and both showed significantly more SGs compared to their corresponding wild-type controls (Figure 3C).

Stress granules can be categorized into canonical and noncanonical categories based on their composition. To define the nature of *Tsc1/2 ^−/−^* induced spontaneous SGs, we characterized their composition by co-staining the SG marker G3BP1 with the canonical translation initiation factors eIF3A, eIF4E, and eIF4G (Figure 3D and Figure 3E). The colocalization observed between G3BP1 and all three factors indicated that the *Tsc1/2 ^−/−^*induced spontaneous SGs are canonical SGs.

Given the observed upregulation of P70 phosphorylation, we hypothesized that the spontaneous SGs were mTORC-1 dependent. To test this hypothesis, we treated *Tsc1/2 ^−/−^*cells with rapamycin, a specific mTORC1 inhibitor, to suppress mTORC1 activity (Figure 3A and Figure 3C). We observed a significant decrease in spontaneous SGs in *Tsc2 ^−/−^* cells, but rapamycin had no effect in *Tsc1 ^−/−^* cells, suggesting *Tsc2 ^−/−^* -induced spontaneous SGs are mTORC1-dependent. The mechanism underlying the rapamycin-insensitivity of spontaneous SGs in *Tsc1 ^−/−^* cells needs further investigation. In summary, these data establish that genetic deletion of *Tsc2* leads to formation of spontaneous and canonical SGs, and this process is sensitive to mTORC1 suppression.

### Spontaneous SGs caused by loss of *Tsc1* and *Tsc2* are translation-dependent

To investigate whether *Tsc1^−/−^* and *Tsc2 ^−/−^* induced spontaneous SGs are in dynamic equilibrium with polysomes, we treated cells with compounds that either promote the release of mRNA from polysomes or confine mRNA into polysomes. We first treated cells with puromycin, a tRNA analog that causes premature termination of translation and the dissociation of polysomes, thus increasing the pool of untranslated mRNA available to aggregate into SGs (Ko et al., 2022, Azzam and Algranati, 1973, Aviner, 2020). Puromycin treatment results in a significant increase in the number of SG-containing cells observed in both *Tsc1^−/−^* and *Tsc2 ^−/−^* cells compared to the corresponding wild type cells (Figure 4A and Figure 4B). This increase in SG formation suggests that *Tsc1^−/−^* and *Tsc2 ^−/−^* induced spontaneous SGs depend on the availability of translational components.

**Figure 4.**
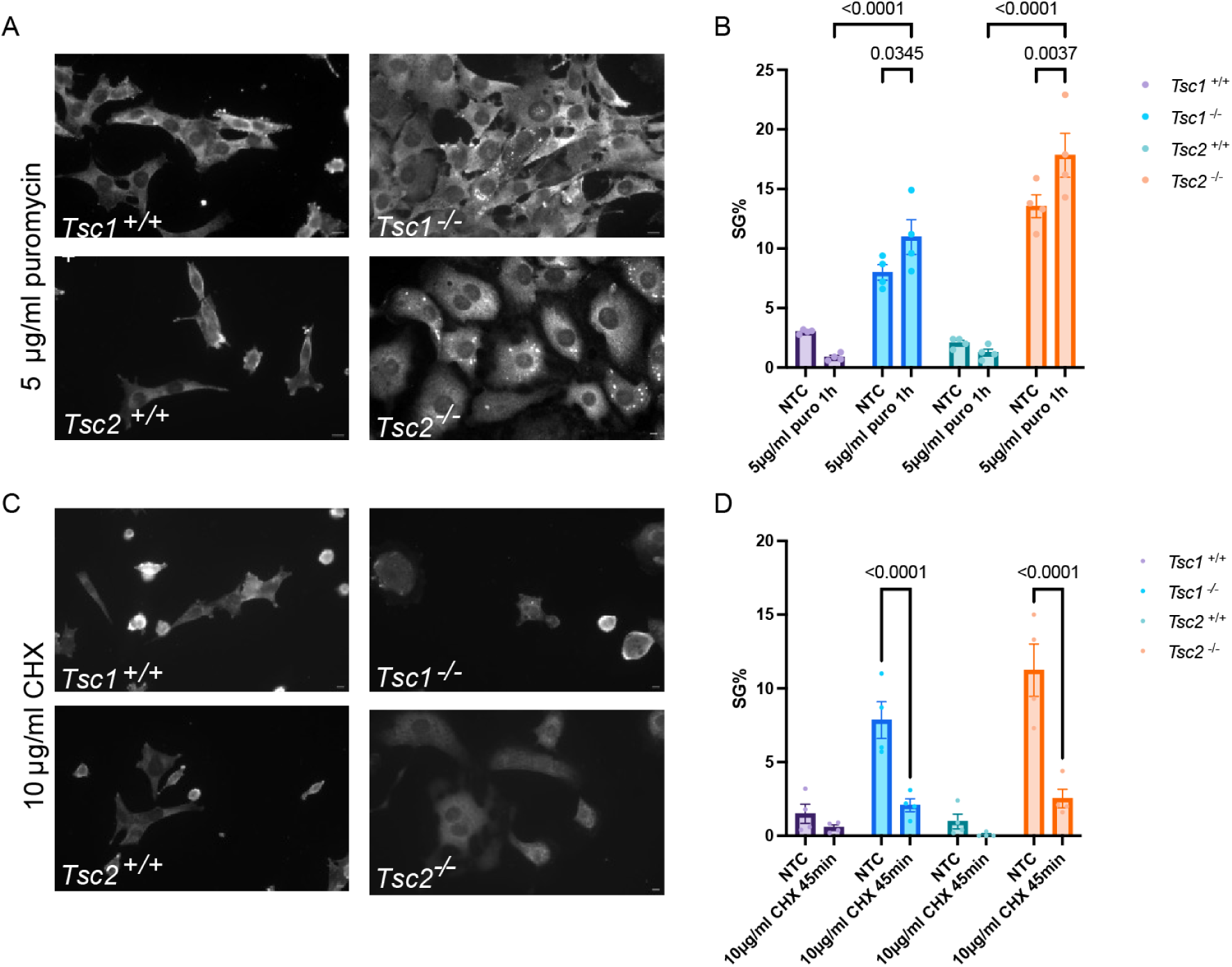
*Tsc1^−/−^* and *Tsc2^−/−^* induced SGs are translation-related. A) Immunofluorescence and B) quantification of SG-positive cells following puromycin treatment. C) Immunofluorescence and D) quantification of SG-positive cells following cycloheximide treatment. Scale bar, 10μm. Data are presented as mean ± SEM of four independent experiments (n=4). Statistical significance was determined using a two-way ANOVA followed by Tukey’s multiple comparisons test.

To further confirm the translation dependency, we treated *Tsc1^−/−^* and *Tsc2 ^−/−^* cells with cycloheximide, an elongation inhibitor that fixes ribosomes onto mRNA, making the mRNA unavailable to aggregate into SGs (Schneider-Poetsch et al., 2010, Santos et al., 2019, Park et al., 2019). As predicted, spontaneous SG formation was significantly suppressed in both *Tsc1^−/−^* and *Tsc2 ^−/−^* cells following cycloheximide treatment (Figure 4C and Figure 4D). In summary, the antagonistic effects of puromycin and cycloheximide confirm that the *Tsc1^−/−^* and *Tsc2 ^−/−^* induced spontaneous SGs are translation-dependent.

### Loss of *Tsc1* or *Tsc2* sensitizes cells to ER stress but not oxidative stress

The loss of *Tsc1* or *Tsc2* leads to hyperactive mTORC1 activity, which consequently drives increased protein synthesis and further overloads the protein folding and processing capacity of the endoplasmic reticulum (ER) (Choi et al., 2018, Qin et al., 2010, Pal et al., 2019, Kosmas et al., 2021, Goncharova et al., 2002). The resulting accumulation of misfolded or unfolded proteins within the ER lumen actives the unfolded protein response (UPR) and further activates the PERK kinase, which subsequently phosphorylates the α-subunit of eukaryotic initiation factor 2 (eIF2α), a key event that inhibits cap-dependent translation initiation and drives SGs formation (Kang et al., 2011).

Given this underlying translational burden, we hypothesized that *Tsc1^−/−^* and *Tsc2 ^−/−^*cells would be hypersensitive to ER stressors. To investigate this sensitivity, we treated *Tsc1^−/−^* and *Tsc2 ^−/−^* cells with two distinct ER stressors known to induce SGs: Thapsigargin (an ER calcium pump inhibitor, Figure 5A and Figure 5B) and MG132 (a proteasome inhibitor that induces ER stress via the accumulation of misfolded protein, Figure 5C and Figure 5D) (Mazroui et al., 2007, Watts et al., 2024, Qin et al., 2010). While all tested cell lines formed SGs following ER stressor treatment, both *Tsc1^−/−^* and *Tsc2 ^−/−^* cells showed a significantly increased percentage of SG-positive cells compared to their corresponding wild type controls, which suggests *Tsc1 or Tsc2* loss lowers the threshold for ER stress-induced SGs formation. Specifically, *Tsc1 ^−/−^* cells exhibited an even greater sensitivity to low doses of Thapsigargin than *Tsc2 ^−/−^* (Figure 5A and Figure 5B), suggesting a unique vulnerability in *Tsc1*^−/−^ cells under mild ER stress.

**Figure 5.**
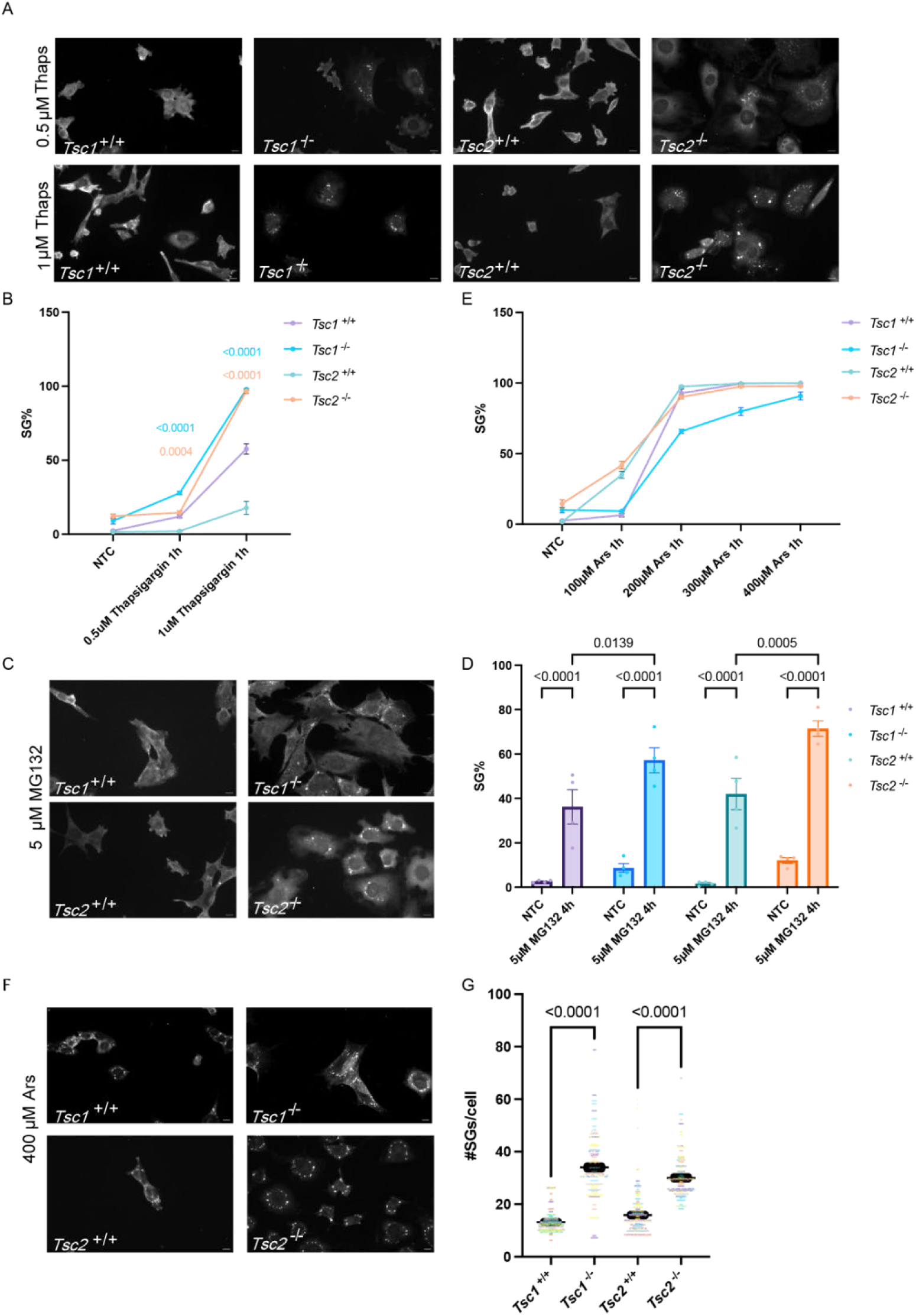
Loss of *Tsc1* or *Tsc2* sensitizes cells to ER stress. A) Immunofluorescence and B) quantification of the percentage of cells with stress granules following Thapsigargin treatment. C) Immunofluorescence and D) quantification of the percentage of cells with stress granules following MG132 treatment. E) Immunofluorescence of SG-positive cells under sodium arsenite treatment. F) Immunofluorescence and G) quantification of the number of SGs per cell per image, following 400uM Ars treatment using CellProfiler. Scale bar, 10 μm. N=4, >120 cells were counted per biological replicate. Data are presented as mean ± SEM of four independent experiments (n=4). Statistical comparisons were performed between the *Tsc1/2 ^−/−^* and *Tsc1/2 ^+/+^* at each concentration using the same color as the line/bar. Statistical significance was determined using a two-way ANOVA followed by Tukey’s multiple comparisons test (B, D, and E) and a one-way ANOVA followed by Šídák’s multiple comparisons test (G).

Given the hypersensitivity to ER stressors, to determine the specificity of this hypersensitivity, we next investigated the impact of *Tsc1* or *Tsc2* loss on the oxidative stress induced by sodium arsenite (Figure 5E). We observed an increase in sensitivity in *Tsc2 ^−/−^*cells compared to *Tsc1 ^−/−^* cells but did not observe any increase in sensitivity when compared to their corresponding control cells. Consistent with the prior findings (Kosmas et al., 2021), we observed a significant increase in number of SGs formed per cell in *Tsc1*^−/−^and *Tsc2*^−/−^ cells (Figure 5F and Figure 5G). This implies that although *Tsc1* or *Tsc2* loss does not lower the threshold for SG formation in response to oxidative stress, the hyperactive mTORC1 activity enhances the cell’s capacity for SGs assembly.

Collectively, *Tsc1* or *Tsc2* deficiency results in a specific hypersensitivity to ER stress, consistent with its role in protein synthesis regulation. The SG response to oxidative stress does not increase in frequency but in magnitude.

### *Tsc1/2 ^−/−^* cells exhibit delayed SG disassembly following ER stress but not oxidative stress

Given that the loss of *Tsc1 or Tsc2* lowers the threshold for ER stress-induced SG assembly, we next hypothesized that loss of *Tsc1 or Tsc2* might also impair the subsequent SG disassembly. To test this hypothesis, we treated cells with ER stressors Thapsigargin (Figure 6A and Figure 6B) and MG132 (Figure 6C and Figure 6D) to induce SG formation, followed by a time course of recovery. Both *Tsc1^−/−^ and Tsc2 ^−/−^* cells lines exhibited a significantly delayed SG disassembly compared to their corresponding wild type controls, suggesting that the loss of *Tsc1 or Tsc2* not only sensitizes cells to ER stressors but also impairs SG disassembly.

**Figure 6.**
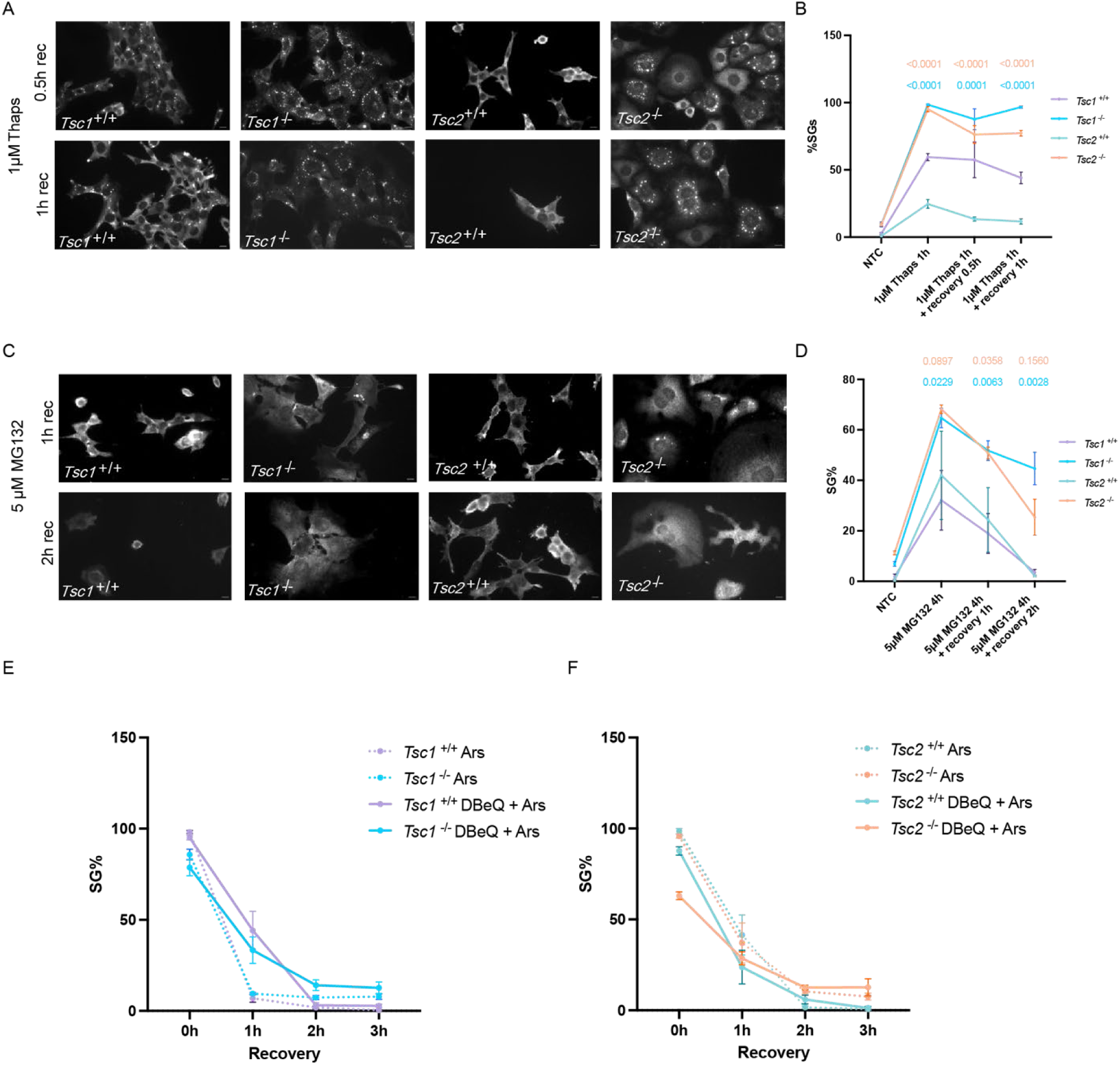
*Tsc1^−/−^* and *Tsc2^−/−^* cells induced SGs have delayed SGs disassembly followed by ER stressors. A) Immunofluorescence and B) quantification of SG-positive cells following Thapsigargin treatment and then allowed to recover for the indicated time points. C) Immunofluorescence and D) quantification of SG-positive cells following MG132 treatment and then allowed to recover for the indicated time points. Quantification of SG-positive cells in E) *Tsc1^−/−^* and F) *Tsc2 ^−/−^ cells* following Ars with and without DBeQ treatment and then allowed to recover for the indicated time points. Scale bar, 10 µM. Data are presented as mean ± SEM of four independent experiments (n=4). Statistical comparisons were performed between the *Tsc1^−/−^ and Tsc2 ^−/−^* cells and corresponding controls cells at each concentration using the same color as the line/bar. Statistical significance was determined using a two-way ANOVA followed by Tukey’s multiple comparisons test.

To determine the specificity of this defect in disassembly, we assessed the disassembly of oxidative stress induced SGs by sodium arsenite (Figure 6E and 6F). Following arsenite treatment and recovery, SG disassembly rate showed no significant difference between *Tsc1^−/−^ and Tsc2 ^−/−^* cells and wild type control cells, which aligns with our previous results (Figure 5C) indicating that the loss of *Tsc1 or Tsc2* does not regulate the threshold for oxidative stress. We further investigated whether inhibition of valosin-containing protein (VCP), an ATPase that plays a role in SG clearance, could affect oxidative stress-induced SG disassembly by co-treating cells with VCP inhibitor DBeQ (Figure 6E and 6F) (Buchan et al., 2013). DBeQ significantly increased SG accumulation in *Tsc2 ^−/−^* cells prior to recovery compared to *Tsc2 ^+/ +^*cells. Following 1h of recovery, we observed significant differences in SG persistence between Ars-DBeQ co-treated *Tsc2 ^−/−^* cells and Ars-treated *Tsc2 ^−/−^* cells, Ars-DBeQ co-treated *Tsc1 ^−/−^*and Ars-treated *Tsc1 ^−/−^*cells, as well as Ars-DBeQ co-treated *Tsc1^+/+^*and Ars-treated *Tsc1^+/+^*cells. Although addition of DBeQ cause SG persistence compared to Ars treatment alone, no significant differential SG disassembly defects between *Tsc1^−/−^ and Tsc2 ^−/−^* cells and corresponding control cells following oxidative stress. This supports the conclusion that the *Tsc1^−/−^ and Tsc2 ^−/−^* cells induced defects in SG disassembly are specific to ER stressors and might not be a general failure to all stress types, even under the additional VCP inhibition.

### Acute *TSC2* depletion sensitizes human cells to ER stress

To determine if the stress granule phenotypes observed in the MEF knockout model are conserved in human cells, we validated our findings in a human model system. We performed transient knockdown of *TSC2* using siRNA in the U2OS cell line. Western blot analysis confirmed high knockdown efficiency, showing a significant reduction in TSC2 protein levels without affecting the TSC1 protein level (Figure 7A and 7B). Consistent with the established function of the TSC complex as a negative regulator of mTORC1, *TSC2* knockdown cells exhibited a statistically significant increase in the ratio of phosphorylated p70 S6 Kinase to total p70 S6 Kinase (Figure 7C). Unlike the *Tsc2^−/−^* MEFs, acute knockdown of *TSC2* in U2OS cells did not result in the robust formation of spontaneous SGs (Figure 7D and 7E).

**Figure 7.**
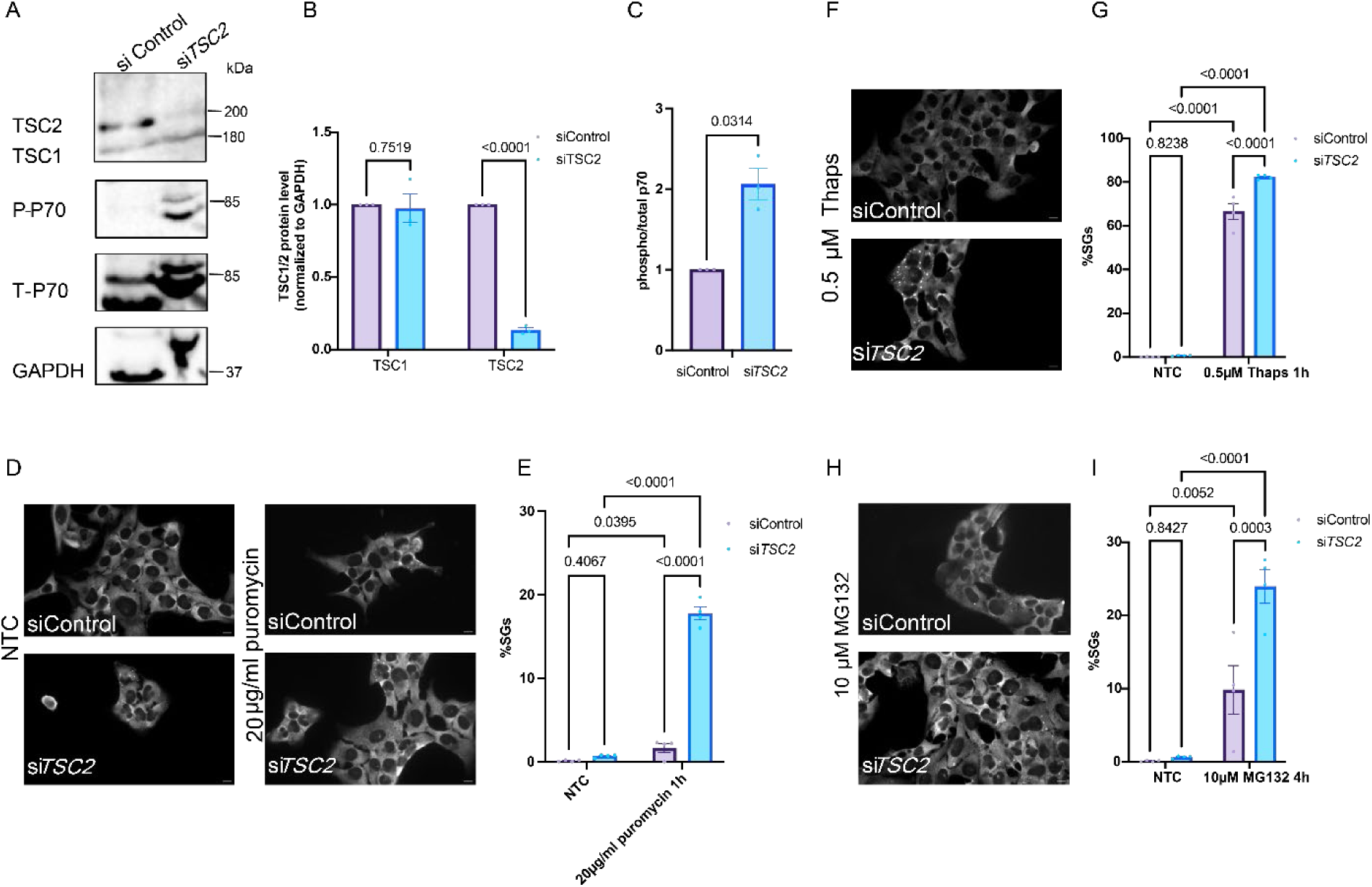
Characterization of *TSC2*-knockdown induced SGs in U2OS cells. A) Western blot analysis of TSC1, TSC2, Phospho-p70 S6 Kinase (P-P70) and Total p70 S6 Kinase (T-P70) in U2OS cells. GAPDH was used as a loading control (mean± SEM, N=3). Quantification of B) TSC1, TSC2, and C) Phospho-p70 S6 Kinase (P-P70) normalized to Total p70 S6 Kinase (T-P70) protein levels ratio in U2OS cells shown in (A). D) Immunofluorescence and E) quantification of the percent of cells with SGs formed in U2OS cells with si*TSC2* were treated with puromycin. F) Immunofluorescence and G) quantification of the percent of cells with SGs formed in U2OS cells with si*TSC2* were treated with Thapsgargin (Thaps). H) Immunofluorescence and I) quantification of the percent of cells with SGs formed in U2OS cells with si*TSC2* were treated with MG132. N=4). Scale bar, 10µM. Statistical significance was determined using E, G, and I) two-way ANOVA with uncorrected Fisher’s LSD and C) unpaired t-test with Welch’s correction.

We next investigated whether cells with *TSC2* knockdown are sensitive to translation manipulation and ER stressors. Upon puromycin treatment, *TSC2* knockdown cells exhibited a significant increase in SG formation compared to control cells (Figure 7D and Figure 7E). *TSC2* knockdown cells also showed a significantly higher percentage of SG-positive cells following treatment with the Thapsigargin (Figure 7F and Figure 7G) and MG132 (Figure 7H and Figure 7I) compared to siControl cells. These findings indicate that while incomplete depletion of *TSC2* suppresses spontaneous SGs formation, the cells are still hypersensitive to translational arrest and ER stress.

### TSC2 does not play a role in pathogenic TDP-43-mediated SG dynamics

SGs have been implicated in pathophysiology of numerous neurodegenerative disorders, most notably Amyotrophic Lateral Sclerosis (ALS), where cytoplasmic inclusions of the RNA-binding protein TAR DNA-binding Protein 43 (TDP-43) represent a pathological hallmark in the majority of cases (Dewey et al., 2012, Streit et al., 2022, Yan et al., 2025, Mori et al., 2024, Dubinski et al., 2023, Niss et al., 2022, Colombrita et al., 2009, Aulas and Vande Velde, 2015). Given the translation-dependent nature of the SGs observed in *Tsc1^−/−^and Tsc2 ^−/−^* MEF models, we tested whether the loss of *TSC2* also exacerbated the progression of pathological TDP-43 aggregates. To test this hypothesis, we co-transfected U2OS cells with an aggregation-prone TDP-43 nuclear localization signal (NLS) mutant alongside siRNA targeting *TSC2* (Figure 8). Previous studies have established that SGs and TDP-43 aggregates undergo a process of mixing and demixing following sodium arsenite treatment (Yan et al., 2025). Our data demonstrated that knocking down *TSC2* did not affect the kinetics of mixing and demixing under the arsenite treatment, suggesting the stress hypersensitivity caused by loss of *TSC2* does not affect pathological TDP-43 aggregates (Figure 8). This suggests that the vulnerability caused by *TSC2* loss is specific to the ER stress-translation axis and does not indiscriminately affect the biophysical properties of pathological aggregates.

**Figure 8.**
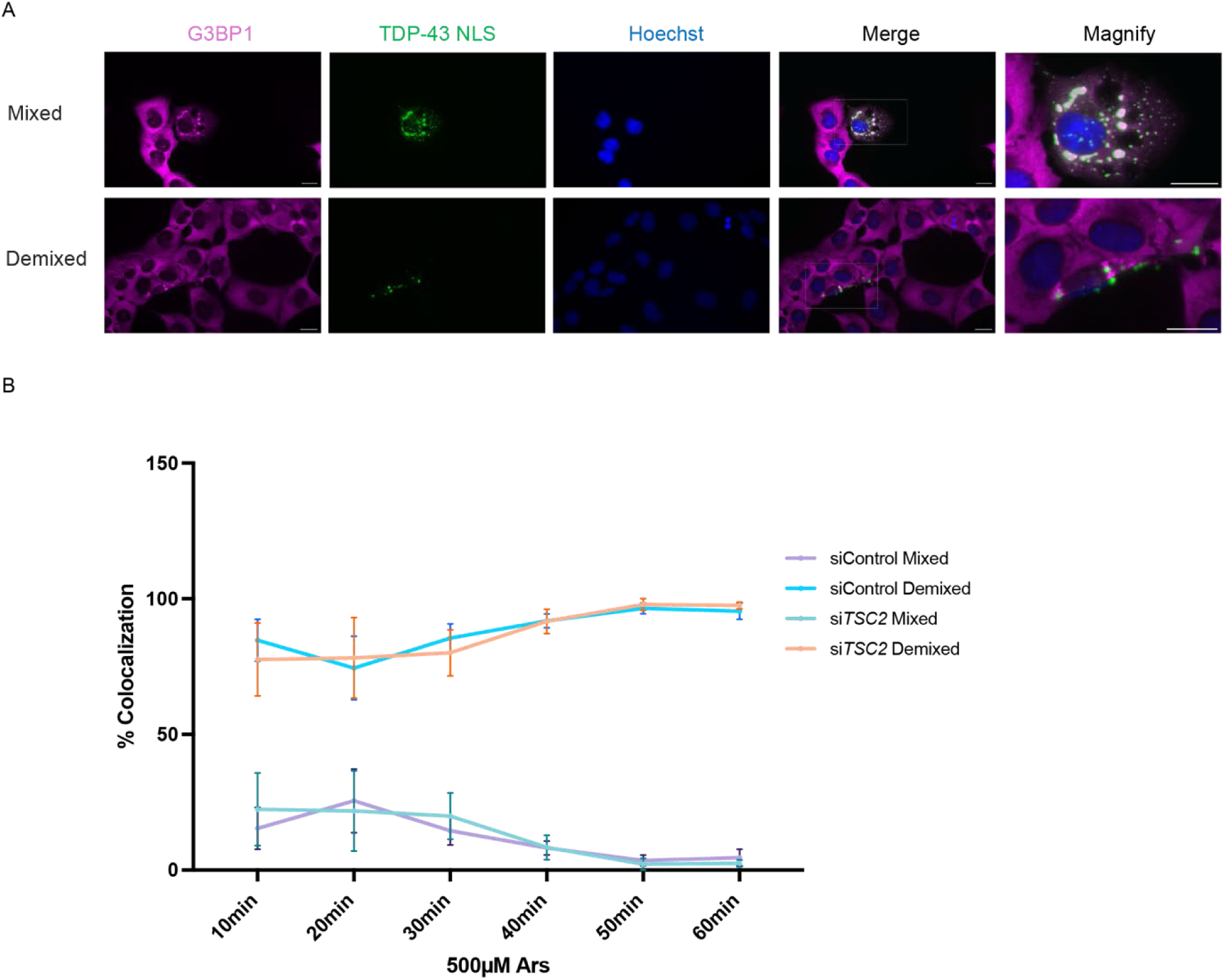
*TSC2* knockdown does not affect TDP-43 NLS aggregation under sodium arsenite stress. A) Representative immunofluorescence images showing the mixed and demixed of G3BP1 and aggregation-prone TDP-43 NLS mutant in U2OS cells. Scale bar, 20 μm. B) Quantification of the colocalization between SGs and TDP-43 NLS in U2OS cells. Cells were co-transfected with an aggregation-prone TDP-43 nuclear localization signal (NLS) mutant and siRNA targeting *TSC2*, followed by treatment with 500 μM Sodium Arsenite for the indicated times. (N=4; over 150 quantified per trial).

### TSC patient derived fibroblast cells maintain normal stress granule homeostasis

To understand the physiological relevance of the phenotype observed in the *Tsc1^−/−^*and *Tsc2 ^−/−^* MEF models, we assessed the stress granule dynamics in tuberous sclerosis complex (TSC) patient derived fibroblast cells (TSP). TSC is a genetic disorder characterized by the widespread growth of benign tumors, driven by *TSC1* or *TSC2* gene mutations and resultant hyperactive mTORC1 signaling. These TSP cells are heterozygous for a large deletion of exons 1 through 14, carrying one null and one functional (wild type) allele.

Western blot analysis confirmed that the TSP cell line showed a reduced basal TSC1 and TSC2 protein level compared to the fibroblast cells derived from an apparently healthy individual (AHI) at the same passage number (Figure 9A). Although the mTORC1 activity, which is calculated by the ratio of phosphorylated p70 S6 kinase and total p70 S6 kinase, appeared upregulated in the TSP cells, this increase was not statistically significant. This is consistent with the lack of spontaneous SGs formation observed in the TSP cells under basal condition (Figure 9B) and their failure to show increased SG assembly following puromycin treatment (Figure 9C).

**Figure 9.**
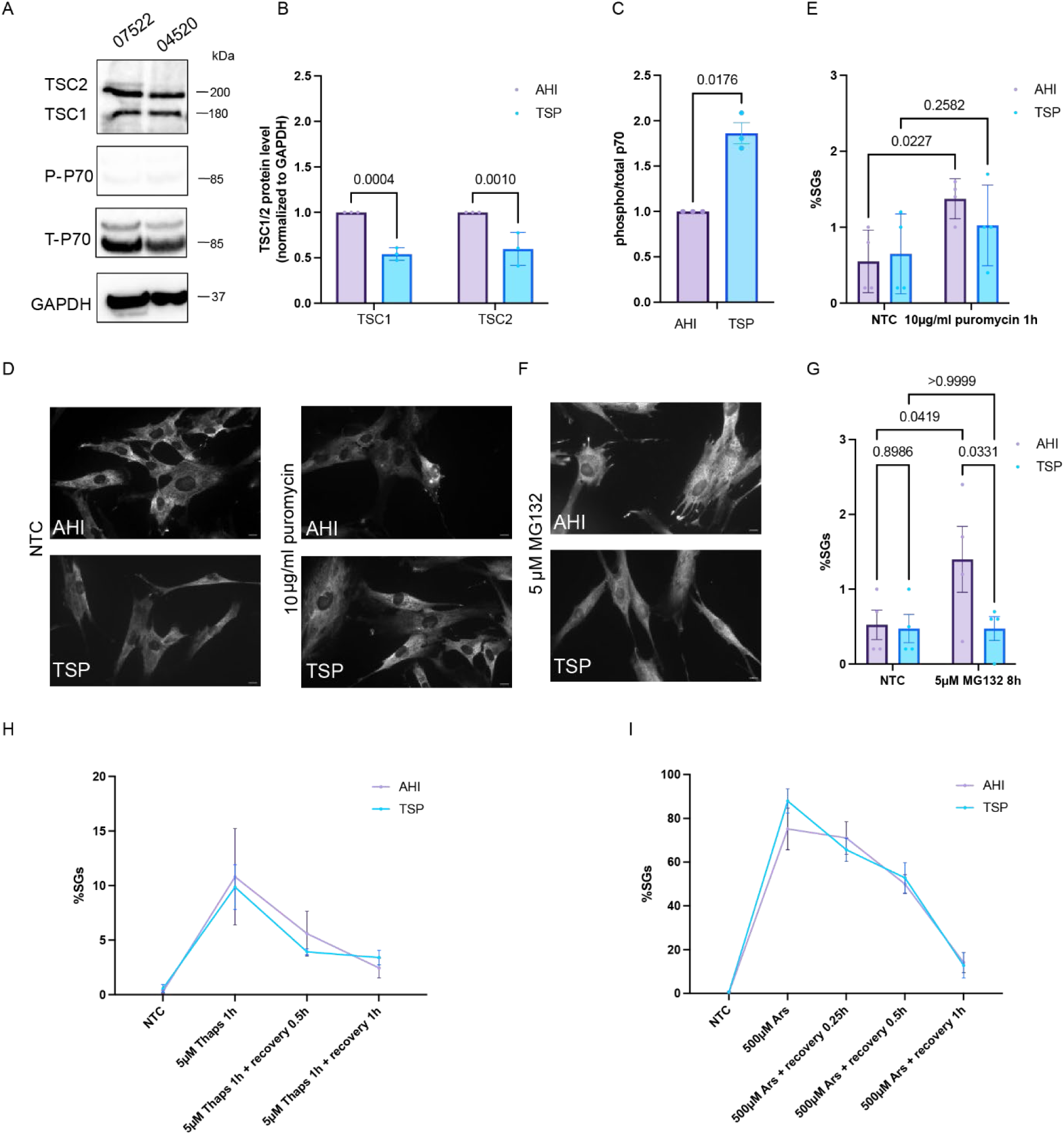
Characterization of fibroblast cells from apparently healthy individual (AHI) and TSC patient (TSP). A) Western blot analysis of TSC1, TSC2, Phospho-p70 S6 Kinase (P-P70) and Total p70 S6 Kinase (T-P70) in fibroblasy cells derived from appreantly healthy individuals (AHI) and TSC patient (TSP). GAPDH was used as a loading control (mean± SEM, N=3). Quantification of B) TSC1, TSC2, and C) Phospho-p70 S6 Kinase (P-P70) normalized to Total p70 S6 Kinase (T-P70) protein levels ratio in AHI and TSP shown in (A). D) Immunofluorescence and E) quantification of percent of cells with SGs formed in AHI and TSP following the puromycin treatment. F) Immunofluorescence and G) quantification of percent of SG-positive cells following the MG132 treatment. Quantification of percent of SGs-positive cells treated with H) Thapsigargin and I) Ars with/without DBeQ pretreatment followed by recovery at indicated times (N=4). Scale bar, 10µM. Data are presented as mean ± SEM. Statistical significance was determined using B, E, and G) two-way ANOVA with uncorrected Fisher’s LSD, C) unpaired t-test with Welch’s correction, and H, I) two-way ANOVA followed by Tukey’s multiple comparisons test.

Next, we investigated whether TSP cells were sensitized to ER stress and oxidative stress. Contradictory to the hypersensitivity observed in the MEF *Tsc1^−/−^ and Tsc2 ^−/−^* cells model, we did not observe any statistical difference between TSP and AHI cells under MG132 (Figure 9F and Figure 9G), Thapsigargin (Figure 9H), and Ars (Figure 9I). Furthermore, SG disassembly kinetics were not impaired in AHI cells following ER stress (Figure 9H) or oxidative stress (Figure 9I). Collectively, these data suggest that the remaining TSC2 present in the human TSC patient fibroblasts is sufficient to suppress the spontaneous SG formation and maintain a normal threshold for SG formation and clearance in response to ER stress and oxidative stress. Greater genetic depletion of *TSC2* is required to create the mTORC1 hyperactivity necessary to disrupt SG homeostasis.

## DISCUSSION

We utilized a high-throughput RNAi screen in *Drosophila* S2R+ cells to identify potential novel SG regulators, revealing *TSC2* as a potent SG suppressor gene. Our approach successfully identified 271 *Drosophila* candidate poly(A)^+^ RNAs granule suppressor genes (Table S1). We hypothesized that part of these poly(A)^+^ RNAs granule suppressor genes identified in *Drosophila* cells could be potential stress granule suppressor genes. Given the link between stress granules and diseases such as neurodegenerative diseases, identifying potential SG suppressor genes on a genome-wide scale could identify possible therapeutic targets for other RNA granule-related diseases and neurodegenerative diseases.

Beyond the priority candidates in Table 1, protein-protein interaction network analysis revealed a significant enrichment of genes that participate in metabolic pathways and growth signaling pathways (Figure S1-S3, Figure 2). A prominent hit within this cluster was the tumor suppressor gene *PTEN*. As a negative regulator of the PI3K/AKT/mTOR signaling pathway, loss of *PTEN* leads to hyperactive mTORC1 activity, promoting SG formation. Du *et al*. reported an increased SG assembly in *PTEN*-deficient cells though both the upregulation of SG nucleator TIAL1 transcription and the activation of PI3K/AKT/mTOR pathway (Du et al., 2024). We also identified *Arm* (human ortholog: *β-Catenin*), which acts as a key nuclear effector of the Wnt signaling pathway and participates in cadherin-based adherens junctions (Vergara et al., 2017). Loss of *β-Catenin* has been shown to reduce mitochondrial mass, which might compromise the ATP production (Vergara et al., 2017, Tang et al., 2022, Jain et al., 2016). This resulting energy stress induces SG formation through the activation of the energy sensor AMPK and the downstream kinase GCN2. Furthermore, the loss of mitochondria mass might lead to additional oxidative stress, which might activate PKR/HRI kinase to drive SG formation (Palma et al., 2024, Girardin et al., 2021). Furthermore, we identified *Su(fu)* (human ortholog: *SUFU*), which is a negative regulator of Hedgehog pathway development (Urman et al., 2016, Katoh and Katoh, 2009). *SUFU* functions at the intersection of Wnt and Hedgehog signaling pathway. Genetic silencing of *SUFU* leads to constitutive activation of Hedgehog pathway which has been shown to upregulate mTORC1 activity. This suggests that the loss of *SUFU* may drive spontaneous SG formation by dysregulated mTORC1 signaling in the similar way as *TSC2* and *PTEN* (Larsen and Møller, 2020, Larsen et al., 2024). The broad association between stress granule formation and signaling pathways suggests stress granule formation is tightly related to cellular fate, translation, and ribosome biogenesis.

We identified 67 candidates with a consistent, robust, spontaneous SG phenotype (Table S2). Among the top ranked genes (Table 1), we selected *TSC2* (*Gig*) as the most promising candidate due to its conserved function within the TSC complex as a negative regulator of mTORC1 (Pal et al., 2019). Because complete loss of *Tsc1* or *Tsc2* is embryonic lethal in mice, we utilized *Tsc1^−/−^* and *Tsc2 ^−/−^* MEF models to validate our screen findings and establish TSC2 as a potent SG suppressor (Ma et al., 2014, Pai et al., 2016, Cormerais et al., 2025, Guijarro et al., 2020). Using *Tsc1^−/−^* and *Tsc2 ^−/−^* cells MEF models, we validated our screen findings and established *TSC2* as a potent SG suppressor gene. Our data confirms that the genetic silencing *Tsc2* leads to hyperactive mTORC1, which is demonstrated by a significant elevated ratio of phosphorylated p70S6K to total p70S6K protein (Figure 3A and Figure 3B). We illustrated that this hyperactivity is the direct driver of spontaneous SG assembly, which is further confirmed by the observation of absence of spontaneous SGs following adding the specific mTORC1 inhibitor rapamycin (Figure 3A and 3C). Interestingly, while *Tsc1^−/−^ and Tsc2 ^−/−^* cells formed spontaneous SGs, rapamycin failed to suppress SG formation in *Tsc1 ^−/−^* cells. This resistance suggests unknown mTORC1-independent functions for TSC1. For instance, loss of *TSC1* has been shown to specifically increase the translation of the nucleolar protein nucleophosmin, which may create a translational burden though an mTORC1-independent mechanism (Mallela and Kumar, 2021). This possibility remains to be fully explored.

We confirm that these spontaneous SGs are translation-dependent, as their formation correlates with the availability of the mRNA pool (Figure 4). Stabilizing polysomes suppressed spontaneous SGs, while dissociating polysomes promoted spontaneous SGs. Previous reports have categorized SGs into canonical and noncanonical categories mainly based on their composition (Cabral et al., 2022). Our immunofluorescence colocalization analysis revealed that *Tsc1^−/−^ and Tsc2 ^−/−^* induced SGs recruit G3BP1 together with translation initiation factors eIF3η, eIF4E, and eIF4G, and they are canonical SGs.

Beyond spontaneous formation, *Tsc2^−/−^* cells have a specific sensitivity to ER stress induced by Thapsigargin and MG132 (Figure 5A and 5B). *Tsc2^−/−^* cells have been shown to display impaired ER stress response, indicated by increased expression of *ATF4*, *ATF6*, and *CHOP* (Kang et al., 2011). Loss of *Tsc2* leads to constitutive activation of mTORC1, which causes elevated translation, suppressed autophagy, and impaired lipid synthesis required for ER membrane expansion (Kang et al., 2011, Johnson et al., 2015, Valvezan and Manning, 2019, Høyer-Hansen and Jäättelä, 2007). The hyperactive mTORC1-induced ER stress could not be rescued by rapamycin, suggesting contributions from rapamycin-insensitive mTOR signaling (Johnson et al., 2015).

In contrast, the response to oxidative stress (sodium arsenite) showed no change in induction threshold but an increase in magnitude of SG formation (Figure 5E-5G). This implies that the loss of *Tsc2* does not universally lower the barrier for all cellular stress. It has been shown that this selective hypersensitivity also appears to other clinically relevant stress pathways. *Tsc1^−/−^ and Tsc2 ^−/−^* cells have been shown to be sensitive to anti-programmed cell death receptor (PD-1) treatment in vivo and transcriptionally upregulated expression of programmed cell death ligand 1 (PD-L1) (Huang et al., 2022). Together, these findings highlight that the dysregulated mTORC1 activity caused by loss of *TSC1* and *TSC2* could be another therapeutic target for treating TSC-associated pathologies.

Relative to SG clearance dynamics, we observed a significant delay in SG disassembly specifically following ER stress (Figure 6A and Figure 6B). SG clearance depends on autophagy and the ubiquitin proteasome system (UPS) (Hu et al., 2022, Wu et al., 2010, Buchan et al., 2013). It is well-established that *Tsc1^−/−^* and *Tsc2 ^−/−^* MEF cells have impaired autophagy initiation and flux (Wu et al., 2010, Choi et al., 2018, Qin et al., 2010, Pal et al., 2019). However, we illustrated that the disassembly kinetics remained normal following oxidative stress, even when the UPS is compromised by addition of the VCP inhibitor DBeQ (Figure 6E and 6F). This suggests that the SG clearance machinery itself may be partially functional but becomes overwhelmed specifically when ER stress generates a massive load of misfolded proteins competing for clearance.

To translate our findings to human models, we utilized U2OS cells and TSC patient-derived fibroblasts (TSP) (Figure 7 and Figure 9). Spontaneous SGs were absent in both *TSC2* knockdown U2OS cells (>95% depletion) and TSP cells (>60% reduction). The TSP cells are heterozygous for a deletion spanning exons 1–14 of *TSC2*, thus western blot analysis still detected approximately 50% of TSC2 protein. Consistent with the embryonic lethality observed in *Tsc2^−/−^* mice (Ma et al., 2014, Pai et al., 2016, Cormerais et al., 2025, Guijarro et al., 2020), these findings suggest that a minimal level of TSC2 is required for organism viability. Notably, the retained C-terminal portion of TSC2 in patient cells may preserve sufficient function to maintain SG homeostasis. However, the high-efficiency knockdown U2OS cells retained specific sensitization to ER stress, whereas the patient fibroblasts did not. Together, these results suggest that complete or near-complete loss of TSC2 (as seen in the null MEFs) is required to drive the spontaneous SG phenotype, while partial reduction or truncation of TSC2 is sufficient to maintain a functional threshold that prevents stress-induced dysregulation.

SGs have been implicated in the pathophysiology of numerous neurodegenerative disorders, especially through their relationship with TDP-43 aggregation. Using an aggregation-prone TDP-43 NLS mutant, we found that loss of *TSC2* did not alter the kinetics of mixing and demixing between TDP-43 aggregates and SGs under oxidative stress, suggesting the hypersensitivity caused by the loss of *TSC2* is specific to ER stress but not affecting other stress or pathological aggregates (Figure 8). However, it is possible that TSC proteins may influence the regulation of other aggregate species under different cellular contexts. Collectively, we define *TSC2* as a critical guardian of SG kinetics, preventing the accumulation of a translational and proteotoxic burden that sensitizes cells to stress. Our screen identified numerous additional candidate SG suppressors, providing targets for future studies to further dissect the signaling pathways that regulate SG dynamics and their connections disease.

## Supporting information

Source Data for Western Blots

## ACKNOWLEDGEMENTS

We thank the Kwiatkowski Laboratory at Dana Farber Cancer Institute for the gift of *Tsc1 -/-*and *Tsc2 -/-* mouse embryonic fibroblasts. N.G.F. was supported by NIH R03AG077140 and R15GM157697, and by new faculty startup funds provided by WPI. Y.M. was supported by the Dr. Armand P. Ferro and Mary H. Ferro Summer Fellowship at WPI.

## AUTHOR CONTRIBUTIONS

Y.M. and N.G.F. conceptualization; Y.M. completed all experiments; Y.M. formal analysis; Y.M. writing—original draft; Y.M. and N.G.F. writing—review & editing; Y.M. Supervision, funding support.

## Online supplementary material

Supplementary Figures and Data accompany this manuscript. Figure S1 shows the protein-protein interaction (PPI) networks of 271 candidate *Drosophila* genes and corresponding GO term analysis with confidence of 0.7, derived from all evidence sources or experimental data only. Figure S2 shows the PPI networks of 271 Drosophila genes and corresponding GO term analysis with confidence of 0.4, derived from all evidence sources. Figure S3 shows the GO term classification from PANTHER classification system. Table S1 lists the combined candidates identified by both normalized and raw foci count approaches yielded 271 genes. Table S2 lists the 67 candidate genes with the most robust and consistent phenotypes across all four replicate images.

## References

Alesi, N., Khabibullin, D., Rosenthal, D. M., Akl, E. W., Cory, P. M., Alchoueiry, M., Salem, S., Daou, M., Gibbons, W. F. & Chen, J. A. 2024. TFEB drives mTORC1 hyperactivation and kidney disease in Tuberous Sclerosis Complex. Nature Communications, 15, 406.

Anderson, P. & Kedersha, N. 2008. Stress granules: the Tao of RNA triage. Trends Biochem Sci, 33, 141–50.

Anderson, P. & Kedersha, N. 2009. Stress granules. Current Biology, 19, R397–R398.

Aulas, A. & Vande Velde, C. 2015. Alterations in stress granule dynamics driven by TDP-43 and FUS: a link to pathological inclusions in ALS? Frontiers in cellular neuroscience, 9, 423.

Aviner, R. 2020. The science of puromycin: From studies of ribosome function to applications in biotechnology. Comput Struct Biotechnol J, 18, 1074–1083.

Azzam, M. & Algranati, I. 1973. Mechanism of puromycin action: fate of ribosomes after release of nascent protein chains from polysomes. Proceedings of the National Academy of Sciences, 70, 3866–3869.

Bakthavachalu, B., Huelsmeier, J., Sudhakaran, I. P., Hillebrand, J., Singh, A., Petrauskas, A., Thiagarajan, D., Sankaranarayanan, M., Mizoue, L., Anderson, E. N., Pandey, U. B., Ross, E., Vijayraghavan, K., Parker, R. & Ramaswami, M. 2018. RNP-Granule Assembly via Ataxin-2 Disordered Domains Is Required for Long-Term Memory and Neurodegeneration. Neuron, 98, 754–766.e4.

Baron, D. M., Kaushansky, L. J., Ward, C. L., Sama, R. R. K., Chian, R.-J., Boggio, K. J., Quaresma, A. J., Nickerson, J. A. & Bosco, D. A. 2013. Amyotrophic lateral sclerosis-linked FUS/TLS alters stress granule assembly and dynamics. Molecular neurodegeneration, 8, 30.

Becker, L. A., Huang, B., Bieri, G., Ma, R., Knowles, D. A., Jafar-Nejad, P., Messing, J., Kim, H. J., Soriano, A., Auburger, G., Pulst, S. M., Taylor, J. P., Rigo, F. & Gitler, A. D. 2017. Therapeutic reduction of ataxin-2 extends lifespan and reduces pathology in TDP-43 mice. Nature, 544, 367–371.

Berchtold, D., Battich, N. & Pelkmans, L. 2018. A Systems-Level Study Reveals Regulators of Membrane-less Organelles in Human Cells. Mol Cell, 72, 1035–1049.e5.

Bjornsti, M.-A. & Houghton, P. J. 2004. The TOR pathway: a target for cancer therapy. Nature Reviews Cancer, 4, 335–348.

Buchan, J. R., Kolaitis, R.-M., Taylor, J. P. & Parker, R. 2013. Eukaryotic Stress Granules Are Cleared by Autophagy and Cdc48/VCP Function. Cell, 153, 1461–1474.

Caban, C., Khan, N., Hasbani, D. M. & Crino, P. B. 2016. Genetics of tuberous sclerosis complex: implications for clinical practice. The application of clinical genetics, 1–8.

Cabral, A. J., Costello, D. C. & Farny, N. G. 2022. The enigma of ultraviolet radiation stress granules: Research challenges and new perspectives. Front Mol Biosci, 9, 1066650.

Cadena Sandoval, M., Heberle, A. M., Rehbein, U., Barile, C., Ramos Pittol, J. M. & Thedieck, K. 2021. mTORC1 crosstalk with stress granules in aging and age-related diseases. Frontiers in aging, 2, 761333.

Chew, J., Cook, C., Gendron, T. F., Jansen-West, K., Del Rosso, G., Daughrity, L. M., Castanedes-Casey, M., Kurti, A., Stankowski, J. N., Disney, M. D., Rothstein, J. D., Dickson, D. W., Fryer, J. D., Zhang, Y. J. & Petrucelli, L. 2019. Aberrant deposition of stress granule-resident proteins linked to C9orf72-associated TDP-43 proteinopathy. Mol Neurodegener, 14, 9.

Chitiprolu, M., Jagow, C., Tremblay, V., Bondy-Chorney, E., Paris, G., Savard, A., Palidwor, G., Barry, F. A., Zinman, L., Keith, J., Rogaeva, E., Robertson, J., Lavallée-Adam, M., Woulfe, J., Couture, J. F., CôTé, J. & Gibbings, D. 2018. A complex of C9ORF72 and p62 uses arginine methylation to eliminate stress granules by autophagy. Nat Commun, 9, 2794.

Choi, H. K., Yuan, H., Fang, F., Wei, X., Liu, L., Li, Q., Guan, J. L. & Liu, F. 2018. Tsc1 regulates the balance between osteoblast and adipocyte differentiation through autophagy/notch1/β-catenin cascade. Journal of Bone and Mineral Research, 33, 2021–2034.

Colombrita, C., Zennaro, E., Fallini, C., Weber, M., Sommacal, A., Buratti, E., Silani, V. & Ratti, A. 2009. TDP-43 is recruited to stress granules in conditions of oxidative insult. Journal of neurochemistry, 111, 1051–1061.

Cormerais, Y., Lapp, S. C., Kalafut, K. C., Cissé, M. Y., Shin, J., Stefadu, B., Personnaz, J., Schrötter, S., Freed, J. & D’amore, A. 2025. AKT-mediated phosphorylation of TSC2 controls stimulus-and tissue-specific mTORC1 signaling and organ growth. Developmental Cell, 60, 2544–2557. e7.

Cui, Q., Liu, Z. & Bai, G. 2024. Friend or foe: The role of stress granule in neurodegenerative disease. Neuron, 112, 2464–2485.

Danos, A. M., Liao, Y., Li, X. & Du, W. 2013. Functional inactivation of Rb sensitizes cancer cells to TSC2 inactivation induced cell death. Cancer letters, 328, 36–43.

De Boer, E. M. J., Orie, V. K., Williams, T., Baker, M. R., De Oliveira, H. M., Polvikoski, T., Silsby, M., Menon, P., Van Den Bos, M. & Halliday, G. M. 2021. TDP-43 proteinopathies: a new wave of neurodegenerative diseases. Journal of Neurology, Neurosurgery & Psychiatry, 92, 86–95.

Dewey, C. M., Cenik, B., Sephton, C. F., Johnson, B. A., Herz, J. & Yu, G. 2012. TDP-43 aggregation in neurodegeneration: Are stress granules the key? Brain Research, 1462, 16–25.

Dibble, C. C., Elis, W., Menon, S., Qin, W., Klekota, J., Asara, J. M., Finan, P. M., Kwiatkowski, D. J., Murphy, L. O. & Manning, B. D. 2012. TBC1D7 is a third subunit of the TSC1-TSC2 complex upstream of mTORC1. Molecular cell, 47, 535–546.

Du, W., Jiang, S., Yin, S., Wang, R., Zhang, C., Yin, B.-C., Li, J., Li, L., Qi, N. & Zhou, Y. 2024. The microbiota-dependent tryptophan metabolite alleviates high-fat diet–induced insulin resistance through the hepatic AhR/TSC2/mTORC1 axis. Proceedings of the National Academy of Sciences, 121, e2400385121.

Dubinski, A., Gagné, M., Peyrard, S., Gordon, D., Talbot, K. & Vande Velde, C. 2023. Stress granule assembly in vivo is deficient in the CNS of mutant TDP-43 ALS mice. Human Molecular Genetics, 32, 319–332.

Farny, N. G., Hurt, J. A. & Silver, P. A. 2008. Definition of global and transcript-specific mRNA export pathways in metazoans. Genes & development, 22, 66–78.

Fernandes, N., Nero, L., Lyons, S. M., Ivanov, P., Mittelmeier, T. M., Bolger, T. A. & Buchan, J. R. 2020. Stress Granule Assembly Can Facilitate but Is Not Required for TDP-43 Cytoplasmic Aggregation. Biomolecules, 10, 1367.

Gan, L., Cookson, M. R., Petrucelli, L. & La Spada, A. R. 2018. Converging pathways in neurodegeneration, from genetics to mechanisms. Nature neuroscience, 21, 1300–1309.

Girardin, S. E., Cuziol, C., Philpott, D. J. & Arnoult, D. 2021. The eIF2α kinase HRI in innate immunity, proteostasis, and mitochondrial stress. The FEBS journal, 288, 3094–3107.

Goncharova, E. A., Goncharov, D. A., Eszterhas, A., Hunter, D. S., Glassberg, M. K., Yeung, R. S., Walker, C. L., Noonan, D., Kwiatkowski, D. J. & Chou, M. M. 2002. Tuberin regulates p70 S6 kinase activation and ribosomal protein S6 phosphorylation: a role for the TSC2 tumor suppressor gene in pulmonary lymphangioleiomyomatosis (LAM). Journal of Biological Chemistry, 277, 30958–30967.

Grao-Cruces, E., Claro-Cala, C. M., Montserrat-De La Paz, S. & Nobrega, C. 2023. Lipoprotein metabolism, protein aggregation, and Alzheimer’s disease: a literature review. International Journal of Molecular Sciences, 24, 2944.

Guijarro, M. V., Danielson, L. S., Cañamero, M., Nawab, A., Abrahan, C., Hernando, E. & Palmer, G. D. 2020. Tsc1 Regulates the proliferation capacity of bone-marrow derived mesenchymal stem cells. Cells, 9, 2072.

Han, L., Wu, Y., Liu, F. & Zhang, H. 2022. eIF4A1 inhibitor suppresses hyperactive mTOR-associated tumors by inducing necroptosis and G2/M arrest. International Journal of Molecular Sciences, 23, 6932.

Haneke, K., Schott, J., Lindner, D., Hollensen, A. K., Damgaard, C. K., Mongis, C., Knop, M., Palm, W., Ruggieri, A. & Stoecklin, G. 2020. CDK1 couples proliferation with protein synthesis. J Cell Biol, 219.

Hans, F., Glasebach, H. & Kahle, P. J. 2020. Multiple distinct pathways lead to hyperubiquitylated insoluble TDP-43 protein independent of its translocation into stress granules. Journal of Biological Chemistry, 295, 673–689.

Hofmann, J. W., Seeley, W. W. & Huang, E. J. 2019. RNA binding proteins and the pathogenesis of frontotemporal lobar degeneration. Annual Review of Pathology: Mechanisms of Disease, 14, 469–495.

Hofmann, S., Kedersha, N., Anderson, P. & Ivanov, P. 2021. Molecular mechanisms of stress granule assembly and disassembly. Biochimica et Biophysica Acta (BBA) - Molecular Cell Research, 1868, 118876.

Høyer-Hansen, M. & Jäättelä, M. 2007. Connecting endoplasmic reticulum stress to autophagy by unfolded protein response and calcium. Cell Death & Differentiation, 14, 1576–1582.

Hu, R., Qian, B., Li, A. & Fang, Y. 2022. Role of proteostasis regulation in the turnover of stress granules. International Journal of Molecular Sciences, 23, 14565.

Huang, H.-S., Chen, L., Chi, J.-X., Lai, S.-Y., Pi, J., Shao, Y.-M. & Xu, J.-F. 2025. Stress granules and cell death: crosstalk and potential therapeutic strategies in infectious diseases. Cell Death & Disease, 16, 495.

Huang, J. & Manning, B. D. 2009. A complex interplay between Akt, TSC2 and the two mTOR complexes. Biochemical Society Transactions, 37, 217–222.

Huang, Q., Li, F., Hu, H., Fang, Z., Gao, Z., Xia, G., Ng, W.-L., Khodadadi-Jamayran, A., Chen, T. & Deng, J. 2022. Loss of TSC1/TSC2 sensitizes immune checkpoint blockade in non–small cell lung cancer. Science advances, 8, eabi9533.

Ishaq, S. M. & Russell, A. P. 2025. Potential role of stress granules and myogranules in amyotrophic lateral sclerosis. Frontiers in Molecular Neuroscience, 18, 1686230.

Jain, S., Wheeler, Joshua R., Walters, Robert W., Agrawal, A., Barsic, A. & Parker, R. 2016. ATPase-Modulated Stress Granules Contain a Diverse Proteome and Substructure. Cell, 164, 487–498.

Johnson, C. E., Hunt, D. K., Wiltshire, M., Herbert, T. P., Sampson, J. R., Errington, R. J., Davies, D. M. & Tee, A. R. 2015. Endoplasmic reticulum stress and cell death in mTORC1-overactive cells is induced by nelfinavir and enhanced by chloroquine. Mol Oncol, 9, 675–88.

Kamagata, K., Iwaki, N., Kanbayashi, S., Banerjee, T., Chiba, R., Gaudon, V., Castaing, B. & Sakomoto, S. 2022. Structure-dependent recruitment and diffusion of guest proteins in liquid droplets of FUS. Scientific reports, 12, 7101.

Kang, Y., Lu, M. & Guan, K. 2011. The TSC1 and TSC2 tumor suppressors are required for proper ER stress response and protect cells from ER stress-induced apoptosis. Cell Death & Differentiation, 18, 133–144.

Karalis, V., Wood, D., Teaney, N. A. & Sahin, M. 2024. The role of TSC1 and TSC2 proteins in neuronal axons. Molecular Psychiatry, 29, 1165–1178.

Katoh, Y. & Katoh, M. 2009. Hedgehog target genes: mechanisms of carcinogenesis induced by aberrant hedgehog signaling activation. Current molecular medicine, 9, 873–886.

Kedersha, N., Cho, M. R., Li, W., Yacono, P. W., Chen, S., Gilks, N., Golan, D. E. & Anderson, P. 2000. Dynamic shuttling of TIA-1 accompanies the recruitment of mRNA to mammalian stress granules. The Journal of cell biology, 151, 1257–1268.

Kedersha, N. L., Gupta, M., Li, W., Miller, I. & Anderson, P. 1999. RNA-binding proteins TIA-1 and TIAR link the phosphorylation of eIF-2α to the assembly of mammalian stress granules. The Journal of cell biology, 147, 1431–1442.

Kim, B., Cooke, H. J. & Rhee, K. 2012. DAZL is essential for stress granule formation implicated in germ cell survival upon heat stress. Development, 139, 568–578.

Ko, T., Oliveira, M. M., Alapin, J. M., Morstein, J., Klann, E. & Trauner, D. 2022. Optical control of translation with a puromycin photoswitch. Journal of the American Chemical Society, 144, 21494–21501.

Kosmas, K., Filippakis, H., Khabibullin, D., Turkiewicz, M., Lam, H. C., Yu, J., Kedersha, N. L., Anderson, P. J. & Henske, E. P. 2021. TSC2 interacts with HDLBP/vigilin and regulates stress granule formation. Molecular Cancer Research, 19, 1389–1397.

Kwiatkowski, D. J., Zhang, H., Bandura, J. L., Heiberger, K. M., Glogauer, M., El-Hashemite, N. & Onda, H. 2002. A mouse model of TSC1 reveals sex-dependent lethality from liver hemangiomas, and up-regulation of p70S6 kinase activity in Tsc1 null cells. Human Molecular Genetics, 11, 525–534.

Larsen, L. J. & Møller, L. B. 2020. Crosstalk of hedgehog and mTORC1 pathways. Cells, 9, 2316.

Larsen, L. J., Østergaard, E. & Møller, L. B. 2024. mTORC1 hampers Hedgehog signaling in Tsc2 deficient cells. Life Science Alliance, 7.

Li, J., Zhang, Y., Gu, J., Zhou, Y., Liu, J., Cui, H., Zhao, T. & Jin, Z. 2024. Stress granule core Protein-Derived peptides inhibit assembly of stress granules and improve Sorafenib sensitivity in cancer cells. Molecules, 29, 2134.

Li, Z., Liu, X. & Liu, M. 2022. Stress granule homeostasis, aberrant phase transition, and amyotrophic lateral sclerosis. ACS Chemical Neuroscience, 13, 2356–2370.

Liu-Yesucevitz, L., Bilgutay, A., Zhang, Y.-J., Vanderwyde, T., Citro, A., Mehta, T., Zaarur, N., Mckee, A., Bowser, R. & Sherman, M. 2010. Tar DNA binding protein-43 (TDP-43) associates with stress granules: analysis of cultured cells and pathological brain tissue. PloS one, 5, e13250.

Luck, K., Kim, D.-K., Lambourne, L., Spirohn, K., Begg, B. E., Bian, W., Brignall, R., Cafarelli, T., Campos-Laborie, F. J., Charloteaux, B., Choi, D., Coté, A. G., Daley, M., Deimling, S., Desbuleux, A., Dricot, A., Gebbia, M., Hardy, M. F., Kishore, N., Knapp, J. J., Kovács, I. A., Lemmens, I., Mee, M. W., Mellor, J. C., Pollis, C., Pons, C., Richardson, A. D., Schlabach, S., Teeking, B., Yadav, A., Babor, M., Balcha, D., Basha, O., Bowman-Colin, C., Chin, S.-F., Choi, S. G., Colabella, C., Coppin, G., D’amata, C., De Ridder, D., De Rouck, S., Duran-Frigola, M., Ennajdaoui, H., Goebels, F., Goehring, L., Gopal, A., Haddad, G., Hatchi, E., Helmy, M., Jacob, Y., Kassa, Y., Landini, S., Li, R., Van Lieshout, N., Macwilliams, A., Markey, D., Paulson, J. N., Rangarajan, S., Rasla, J., Rayhan, A., Rolland, T., San-Miguel, A., Shen, Y., Sheykhkarimli, D., Sheynkman, G. M., Simonovsky, E., Taşan, M., Tejeda, A., Tropepe, V., Twizere, J.-C., Wang, Y., Weatheritt, R. J., Weile, J., Xia, Y., Yang, X., Yeger-Lotem, E., Zhong, Q., Aloy, P., Bader, G. D., De Las Rivas, J., Gaudet, S., Hao, T., Rak, J., Tavernier, J., Hill, D. E., Vidal, M., Roth, F. P. & Calderwood, M. A. 2020. A reference map of the human binary protein interactome. Nature, 580, 402–408.

Ma, A., Wang, L., Gao, Y., Chang, Z., Peng, H., Zeng, N., Gui, Y.-S., Tian, X., Li, X. & Cai, B. 2014. Tsc1 deficiency-mediated mTOR hyperactivation in vascular endothelial cells causes angiogenesis defects and embryonic lethality. Human Molecular Genetics, 23, 693–705.

Ma, Y. & Farny, N. G. 2023. Connecting the dots: Neuronal senescence, stress granules, and neurodegeneration. Gene, 871, 147437.

Maharjan, N., Künzli, C., Buthey, K. & Saxena, S. 2017. C9ORF72 Regulates Stress Granule Formation and Its Deficiency Impairs Stress Granule Assembly, Hypersensitizing Cells to Stress. Mol Neurobiol, 54, 3062–3077.

Mallela, K. & Kumar, A. 2021. Role of TSC1 in physiology and diseases. Molecular and cellular biochemistry, 476, 2269–2282.

Marcelo, A., Koppenol, R., De Almeida, L. P., Matos, C. A. & Nóbrega, C. 2021. Stress granules, RNA-binding proteins and polyglutamine diseases: too much aggregation? Cell Death Dis, 12, 592.

Marqués, P., Burillo, J., González-Blanco, C., Jiménez, B., García, G., García-Aguilar, A., Iglesias-Fortes, S., Lockwood, Á. & Guillén, C. 2024. Regulation of TSC2 lysosome translocation and mitochondrial turnover by TSC2 acetylation status. Scientific Reports, 14, 12521.

Mazroui, R., Di Marco, S., Kaufman, R. J. & Gallouzi, I.-E. 2007. Inhibition of the ubiquitin-proteasome system induces stress granule formation. Molecular biology of the cell, 18, 2603–2618.

Mccubrey, J. A., Steelman, L. S., Chappell, W. H., Abrams, S. L., Montalto, G., Cervello, M., Nicoletti, F., Fagone, P., Malaponte, G., Mazzarino, M. C., Candido, S., Libra, M., Bäsecke, J., Mijatovic, S., Maksimovic-Ivanic, D., Milella, M., Tafuri, A., Cocco, L., Evangelisti, C., Chiarini, F. & Martelli, A. M. 2012. Mutations and deregulation of Ras/Raf/MEK/ERK and PI3K/PTEN/Akt/mTOR cascades which alter therapy response. Oncotarget, 3, 954–87.

Mcdonald, K. K., Aulas, A., Destroismaisons, L., Pickles, S., Beleac, E., Camu, W., Rouleau, G. A. & Vande Velde, C. 2011. TAR DNA-binding protein 43 (TDP-43) regulates stress granule dynamics via differential regulation of G3BP and TIA-1. Human Molecular Genetics, 20, 1400–1410.

Millar, S. R., Huang, J. Q., Schreiber, K. J., Tsai, Y.-C., Won, J., Zhang, J., Moses, A. M. & Youn, J.-Y. 2023. A new phase of networking: the molecular composition and regulatory dynamics of mammalian stress granules. Chemical reviews, 123, 9036–9064.

Mokas, S., Mills, J. R., Garreau, C., Fournier, M.-J., Robert, F., Arya, P., Kaufman, R. J., Pelletier, J. & Mazroui, R. 2009. Uncoupling stress granule assembly and translation initiation inhibition. Molecular biology of the cell, 20, 2673–2683.

Mori, F., Yasui, H., Miki, Y., Kon, T., Arai, A., Kurotaki, H., Tomiyama, M. & Wakabayashi, K. 2024. Colocalization of TDP-43 and stress granules at the early stage of TDP-43 aggregation in amyotrophic lateral sclerosis. Brain Pathology, 34, e13215.

Nahm, M., Lim, S. M., Kim, Y. E., Park, J., Noh, M. Y., Lee, S., Roh, J. E., Hwang, S. M., Park, C. K., Kim, Y. H., Lim, G., Lee, J., Oh, K. W., Ki, C. S. & Kim, S. H. 2020. ANXA11 mutations in ALS cause dysregulation of calcium homeostasis and stress granule dynamics. Sci Transl Med, 12.

Nedelsky, N. B. & Taylor, J. P. 2022. Pathological phase transitions in ALS-FTD impair dynamic RNA–protein granules. Rna, 28, 97–113.

Niss, F., Piñero-Paez, L., Zaidi, W., Hallberg, E. & Ström, A.-L. 2022. Key modulators of the stress granule response TIA1, TDP-43, and G3BP1 are altered by polyglutamine-expanded ATXN7. Molecular Neurobiology, 59, 5236–5251.

Ohn, T., Kedersha, N., Hickman, T., Tisdale, S. & Anderson, P. 2008. A functional RNAi screen links O-GlcNAc modification of ribosomal proteins to stress granule and processing body assembly. Nature cell biology, 10, 1224–1231.

Onda, H., Lueck, A., Marks, P. W., Warren, H. B. & Kwiatkowski, D. J. 1999. Tsc2+/– mice develop tumors in multiple sites that express gelsolin and are influenced by genetic background. The Journal of clinical investigation, 104, 687–695.

Pai, G. M., Zielinski, A., Koalick, D., Ludwig, K., Wang, Z.-Q., Borgmann, K., Pospiech, H. & Rubio, I. 2016. TSC loss distorts DNA replication programme and sensitises cells to genotoxic stress. Oncotarget, 7, 85365.

Pal, R., Xiong, Y. & Sardiello, M. 2019. Abnormal glycogen storage in tuberous sclerosis complex caused by impairment of mTORC1-dependent and-independent signaling pathways. Proceedings of the National Academy of Sciences, 116, 2977–2986.

Palma, F. R., Gantner, B. N., Sakiyama, M. J., Kayzuka, C., Shukla, S., Lacchini, R., Cunniff, B. & Bonini, M. G. 2024. ROS production by mitochondria: function or dysfunction? Oncogene, 43, 295–303.

Park, N. Y., Heo, Y., Yang, J. W., Yoo, J. M., Jang, H. J., Jo, J. H., Park, S. J., Lin, Y., Choi, J. & Jeon, H. 2025. Graphene quantum dots attenuate TDP-43 proteinopathy in amyotrophic lateral sclerosis. Acs Nano, 19, 8692–8710.

Park, Y., Koga, Y., Su, C., Waterbury, A. L., Johnny, C. L. & Liau, B. B. 2019. Versatile synthetic route to cycloheximide and analogues that potently inhibit translation elongation. Angewandte Chemie, 131, 5441–5445.

Parobkova, E. & Matej, R. 2021. Amyotrophic Lateral Sclerosis and Frontotemporal Lobar Degenerations: Similarities in Genetic Background. Diagnostics (Basel), 11.

Patursky-Polischuk, I., Stolovich-Rain, M., Hausner-Hanochi, M., Kasir, J., Cybulski, N., Avruch, J., Rüegg, M. A., Hall, M. N. & Meyuhas, O. 2009. The TSC-mTOR pathway mediates translational activation of TOP mRNAs by insulin largely in a raptor-or rictor-independent manner. Molecular and cellular biology.

Polymenidou, M. & Cleveland, D. W. 2011. The seeds of neurodegeneration: prion-like spreading in ALS. Cell, 147, 498–508.

Potter, C. J., Pedraza, L. G. & Xu, T. 2002. Akt regulates growth by directly phosphorylating Tsc2. Nature cell biology, 4, 658–665.

Qin, J., Wang, Z., Hoogeveen-Westerveld, M., Shen, G., Gong, W., Nellist, M. & Xu, W. 2016. Structural basis of the interaction between tuberous sclerosis complex 1 (TSC1) and Tre2-Bub2-Cdc16 domain family member 7 (TBC1D7). Journal of Biological Chemistry, 291, 8591–8601.

Qin, L., Wang, Z., Tao, L. & Wang, Y. 2010. ER stress negatively regulates AKT/TSC/mTOR pathway to enhance autophagy. Autophagy, 6, 239–247.

Ramaswami, M., Taylor, J. P. & Parker, R. 2013. Altered ribostasis: RNA-protein granules in degenerative disorders. Cell, 154, 727–736.

Reyna-Fabián, M. E., Hernández-Martínez, N. L., Alcántara-Ortigoza, M. A., Ayala-Sumuano, J. T., Enríquez-Flores, S., Velázquez-Aragón, J. A., Varela-Echavarría, A., Todd-Quiñones, C. G. & González-Del Angel, A. 2020. First comprehensive TSC1/TSC2 mutational analysis in Mexican patients with Tuberous Sclerosis Complex reveals numerous novel pathogenic variants. Scientific reports, 10, 6589.

Rosset, C., Netto, C. B. O. & Ashton-Prolla, P. 2017. TSC1 and TSC2 gene mutations and their implications for treatment in Tuberous Sclerosis Complex: a review. Genetics and molecular biology, 40, 69–79.

Rummens, J., Khalil, B., Yıldırım, G., Silva, P., Zorzini, V., Peredo, N., Wojno, M., Ramakers, M., Van Den Bosch, L., Van Damme, P., Davie, K., Hendrix, J., Rousseau, F., Schymkowitz, J. & Da Cruz, S. 2025. TDP-43 seeding induces cytoplasmic aggregation heterogeneity and nuclear loss of function of TDP-43. Neuron, 113, 1597–1613.e8.

Santiago Lima, A. J., Hoogeveen-Westerveld, M., Nakashima, A., Maat-Kievit, A., Van Den Ouweland, A., Halley, D., Kikkawa, U. & Nellist, M. 2014. Identification of regions critical for the integrity of the TSC1-TSC2-TBC1D7 complex. PLoS One, 9, e93940.

Santos, D. A., Shi, L., Tu, B. P. & Weissman, J. S. 2019. Cycloheximide can distort measurements of mRNA levels and translation efficiency. Nucleic acids research, 47, 4974–4985.

Schneider-Poetsch, T., Ju, J., Eyler, D. E., Dang, Y., Bhat, S., Merrick, W. C., Green, R., Shen, B. & Liu, J. O. 2010. Inhibition of eukaryotic translation elongation by cycloheximide and lactimidomycin. Nature chemical biology, 6, 209–217.

Scialò, C., Zhong, W., Jagannath, S., Wilkins, O., Caredio, D., Hruska-Plochan, M., Lurati, F., Peter, M., De Cecco, E., Celauro, L., Aguzzi, A., Legname, G., Fratta, P. & Polymenidou, M. 2025. Seeded aggregation of TDP-43 induces its loss of function and reveals early pathological signatures. Neuron, 113, 1614–1628.e11.

Seguin, S. J., Morelli, F. F., Vinet, J., Amore, D., De Biasi, S., Poletti, A., Rubinsztein, D. C. & Carra, S. 2014. Inhibition of autophagy, lysosome and VCP function impairs stress granule assembly. Cell Death Differ, 21, 1838–51.

Shannon, P., Markiel, A., Ozier, O., Baliga, N. S., Wang, J. T., Ramage, D., Amin, N., Schwikowski, B. & Ideker, T. 2003. Cytoscape: a software environment for integrated models of biomolecular interaction networks. Genome Res, 13, 2498–504.

Smith, J. & Bartel, D. P. 2026. The G3BP stress-granule proteins reinforce the integrated stress response translation programme. Nature Cell Biology, 28, 135–148.

Streit, L., Kuhn, T., Vomhof, T., Bopp, V., Ludolph, A. C., Weishaupt, J. H., Gebhardt, J. C. M., Michaelis, J. & Danzer, K. M. 2022. Stress induced TDP-43 mobility loss independent of stress granules. Nature Communications, 13, 5480.

Sun, Y., Yang, P., Zhang, Y., Bao, X., Li, J., Hou, W., Yao, X., Han, J. & Zhang, H. 2011. A genome-wide RNAi screen identifies genes regulating the formation of P bodies in C. elegans and their functions in NMD and RNAi. Protein & Cell, 2, 918–939.

Szklarczyk, D., Kirsch, R., Koutrouli, M., Nastou, K., Mehryary, F., Hachilif, R., Gable, A. L., Fang, T., Doncheva, N. T., Pyysalo, S., Bork, P., Jensen, L. J. & Von Mering, C. 2023. The STRING database in 2023: protein-protein association networks and functional enrichment analyses for any sequenced genome of interest. Nucleic Acids Res, 51, D638–d646.

Tang, C. P., Clark, O., Ferrarone, J. R., Campos, C., Lalani, A. S., Chodera, J. D., Intlekofer, A. M., Elemento, O. & Mellinghoff, I. K. 2022. GCN2 kinase activation by ATP-competitive kinase inhibitors. Nature chemical biology, 18, 207–215.

Ueda, T., Takeuchi, T., Fujikake, N., Suzuki, M., Minakawa, E. N., Ueyama, M., Fujino, Y., Kimura, N., Nagano, S. & Yokoseki, A. 2024. Dysregulation of stress granule dynamics by DCTN1 deficiency exacerbates TDP-43 pathology in Drosophila models of ALS/FTD. Acta Neuropathologica Communications, 12, 20.

Urman, N. M., Mirza, A., Atwood, S. X., Whitson, R. J., Sarin, K. Y., Tang, J. Y. & Oro, A. E. 2016. Tumor-Derived Suppressor of Fused Mutations Reveal Hedgehog Pathway Interactions. PLoS One, 11, e0168031.

Valvezan, A. J. & Manning, B. D. 2019. Molecular logic of mTORC1 signalling as a metabolic rheostat. Nat Metab, 1, 321–333.

Vergara, D., Stanca, E., Guerra, F., Priore, P., Gaballo, A., Franck, J., Simeone, P., Trerotola, M., De Domenico, S., Fournier, I., Bucci, C., Salzet, M., Giudetti, A. M. & Maffia, M. 2017. β-Catenin Knockdown Affects Mitochondrial Biogenesis and Lipid Metabolism in Breast Cancer Cells. Front Physiol, 8, 544.

Wang, K., Lockwood, S. E. & Manning, B. D. 2025a. Evolution of growth factor signaling to the TSC complex to regulate mTORC1. Science Signaling, 18, eadw4165.

Wang, S., Ma, R., Gao, C., Tian, Y.-N., Hu, R.-G., Zhang, H., Li, L. & Li, Y. 2025b. Unraveling the function of TSC1-TSC2 complex: implications for stem cell fate. Stem Cell Research & Therapy, 16, 38.

Watts, M. E., Giadone, R. M., Ordureau, A., Holton, K. M., Harper, J. W. & Rubin, L. L. 2024. Analyzing the ER stress response in ALS patient derived motor neurons identifies druggable neuroprotective targets. Frontiers in Cellular Neuroscience, 17, 1327361.

Wheeler, E. C., Vu, A. Q., Einstein, J. M., Disalvo, M., Ahmed, N., Van Nostrand, E. L., Shishkin, A. A., Jin, W., Allbritton, N. L. & Yeo, G. W. 2020. Pooled CRISPR screens with imaging on microraft arrays reveals stress granule-regulatory factors. Nature methods, 17, 636–642.

White, J. P. & Lloyd, R. E. 2011. Poliovirus unlinks TIA1 aggregation and mRNA stress granule formation. Journal of virology, 85, 12442–12454.

Wu, Y., Wang, X., Meng, L., Liao, Z., Ji, W., Zhang, P., Lin, J. & Guo, Q. 2025. Translation landscape of stress granules. Science Advances, 11, eady6859.

Wu, Y.-T., Tan, H.-L., Shui, G., Bauvy, C., Huang, Q., Wenk, M. R., Ong, C.-N., Codogno, P. & Shen, H.-M. 2010. Dual role of 3-methyladenine in modulation of autophagy via different temporal patterns of inhibition on class I and III phosphoinositide 3-kinase. Journal of Biological Chemistry, 285, 10850–10861.

Xing, F., Qin, Y., Xu, J., Wang, W. & Zhang, B. 2023. Stress granules dynamics and promising functions in pancreatic cancer. Biochimica et Biophysica Acta (BBA)-Reviews on Cancer, 1878, 188885.

Yan, X., Kuster, D., Mohanty, P., Nijssen, J., Pombo-García, K., Garcia Morato, J., Rizuan, A., Franzmann, T. M., Sergeeva, A., Ly, A. M., Liu, F., Passos, P. M., George, L., Wang, S.-H., Shenoy, J., Danielson, H. L., Ozguney, B., Honigmann, A., Ayala, Y. M., Fawzi, N. L., Dickson, D. W., Rossoll, W., Mittal, J., Alberti, S. & Hyman, A. A. 2025. Intra-condensate demixing of TDP-43 inside stress granules generates pathological aggregates. Cell, 188, 4123–4140.e18.

Yang, P., Mathieu, C., Kolaitis, R.-M., Zhang, P., Messing, J., Yurtsever, U., Yang, Z., Wu, J., Li, Y. & Pan, Q. 2020. G3BP1 is a tunable switch that triggers phase separation to assemble stress granules. Cell, 181, 325–345. e28.

Youn, J.-Y., Dunham, W. H., Hong, S. J., Knight, J. D., Bashkurov, M., Chen, G. I., Bagci, H., Rathod, B., Macleod, G. & Eng, S. W. 2018. High-density proximity mapping reveals the subcellular organization of mRNA-associated granules and bodies. Molecular cell, 69, 517–532. e11.

Yuan, L., Mao, L. H., Huang, Y. Y., Outeiro, T. F., Li, W., Vieira, T. & Li, J. Y. 2025. Stress granules: emerging players in neurodegenerative diseases. Transl Neurodegener, 14, 22.

Yuan, X., Wang, Q., Dai, M., Wang, H., Xiong, X., Pan, L. & Wang, C. 2024. TSC2 gene characterization and mechanism of ammonia nitrogen stress inhibiting growth through AMPK/mTOR pathway mediated by TSC2 in Megalobrama amblycephala. Aquaculture Reports, 38, 102342.

Zhang, H., Cicchetti, G., Onda, H., Koon, H. B., Asrican, K., Bajraszewski, N., Vazquez, F., Carpenter, C. L. & Kwiatkowski, D. J. 2003. Loss of Tsc1/Tsc2 activates mTOR and disrupts PI3K-Akt signaling through downregulation of PDGFR. The Journal of clinical investigation, 112, 1223–1233.

Zheng, W., Wang, K., Wu, Y., Yan, G., Zhang, C., Li, Z., Wang, L. & Chen, S. 2022. C9orf72 regulates the unfolded protein response and stress granule formation by interacting with eIF2α. Theranostics, 12, 7289–7306.

Zhu, Q.-Y., He, Z.-M., Cao, W.-M. & Li, B. 2023. The role of TSC2 in breast cancer: a literature review. Frontiers in Oncology, 13, 1188371.

