## Supplementary Information for "TSC2 is a stress granule suppressor"

The supplementary information accompanying this manuscript consists of three supplementary figures (in this file) and two supplementary tables (in an additional Excel file).

A

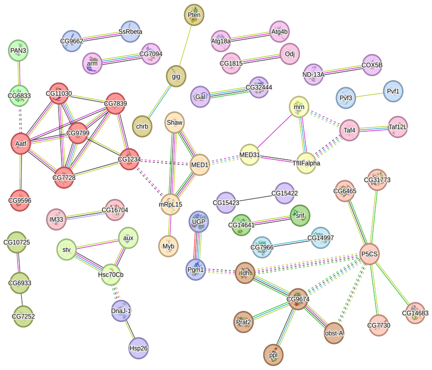


B

| Color | Cluster ID | Gene Count | Description |
|---|---|---|---|
| | Cluster 1 | 8 | Preribosome and Nucleolus |
| | Cluster 2 | 3 | Basal transcription factors |
| | Cluster 3 | 3 | Chitin binding domain superfamily, and Protein TsetseEP |
| | Cluster 4 | 3 | Stress response |
| | Cluster 5 | 3 | FBgn0003386, FBgn0036990, FBgn0037109 |
| | Cluster 6 | 2 | snRNA binding |
| | Cluster 7 | 2 | Oligosaccharyltransferase complex, and Post-translational protein targeting to membrane, translocation |
| | Cluster 8 | 2 | FBgn031097, FBgn0036663 |
| | Cluster 9 | 2 | Transcription factor TFIID complex |
| | Cluster 10 | 2 | Mixed, incl. Aspartyl protease, and Calycin |
| | Cluster 11 | 2 | FBgn0031684, FBgn0031830 |
| | Cluster 12 | 2 | BPTI/Kunitz family of serine protease inhibitors |

C

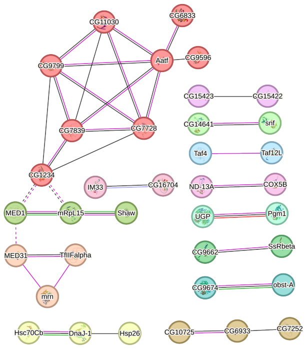


D

| Color | Cluster ID | Gene Count | Description |
|---|---|---|---|
| | Cluster 1 | 8 | Preribosome, and Nucleolus |
| | Cluster 2 | 3 | 1. Basal transcription factors<br>2. RNA Pol II CTD phosphorylation and interaction with CE |
| | Cluster 3 | 3 | Chitin binding domain superfamily, and Protein TsetseEP |
| | Cluster 4 | 3 | 1. Stress response<br>2. Hsp20/alpha crystallin family, and Heat shock protein 70, conserved site |
| | Cluster 5 | 3 | MED1, Shaw, mRpL15 |
| | Cluster 6 | 2 | snRNA binding |
| | Cluster 7 | 2 | Oligosaccharyltransferase complex, and Post-translational protein |
| | Cluster 8 | 2 | 1. Glycogen synthesis<br>2. Starch and sucrose metabolism |
| | Cluster 9 | 2 | CG9674, obst-A |
| | Cluster 10 | 2 | Transcription factor TFIID complex |
| | Cluster 11 | 2 | Mixed, incl. Aspartyl protease, and Calycin |
| | Cluster 12 | 2 | COX5B, ND-13A |
| | Cluster 13 | 2 | 1. BPTI/Kunitz family of serine protease inhibitors<br>2. Mixed, incl. Transforming growth factor beta binding, and Lipoprotein transport |

**Figure S1 Comparative Interactome Analysis of Identified *Drosophila* Stress Granule Suppressors**

A) Protein–protein interaction (PPI) analysis of the 271 candidate *Drosophila* genes identified from the RNAi re-analysis was performed using all evidence sources from STRING, including text mining, experiments, databases, co-expression, neighborhood, gene fusion, and co-occurrence. The network was filtered with a confidence score of 0.7. B) Major functional clusters are color-coded according to the Gene Ontology (GO) terms identified in the enrichment analysis. C) PPI network of the same 271 *Drosophila* genes using a restricted evidence set, including experiments, co-expression, neighborhood, gene fusion, and co-occurrence. The network was filtered with a confidence score of 0.7. D) Major functional clusters are color-coded according to the Gene Ontology (GO) terms identified in the enrichment analysis.

A

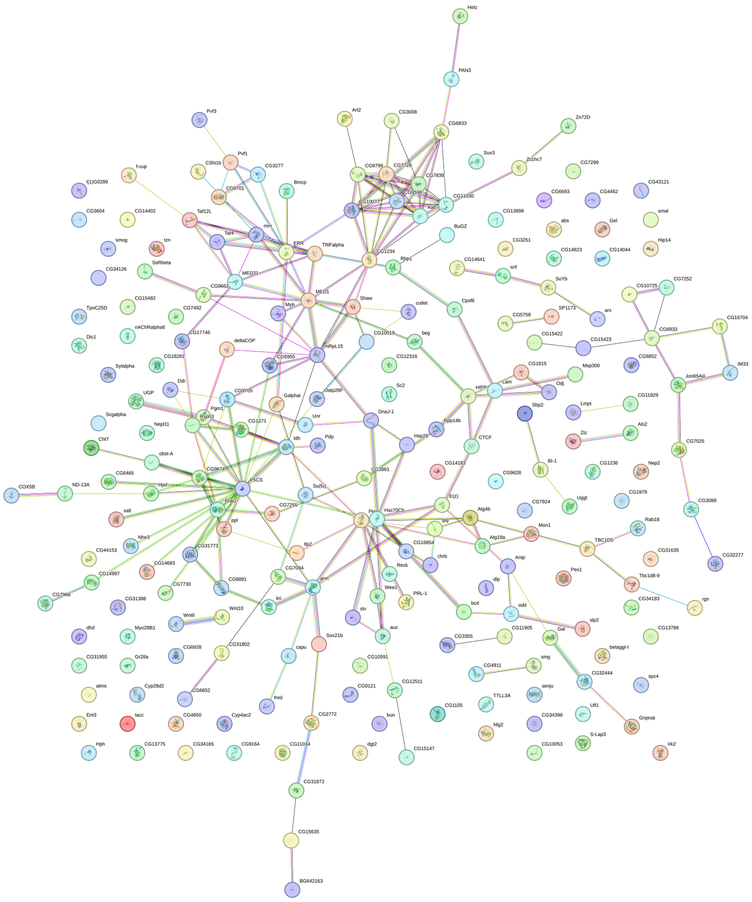

B

**KEGG Pathways**

| Pathway ID | Description | Count in Network | Strength | Signal | False Discovery Rate |
|---|---|---|---|---|---|
| dme04310 | Wnt signaling pathway | 6 of 102 | 1.32 | 1.23 | 6.60E-05 |
| dme00052 | Galactose metabolism | 3 of 26 | 1.47 | 0.68 | 0.0115 |
| dme04150 | mTOR signaling pathway | 4 of 99 | 1.16 | 0.62 | 0.0115 |
| dme01230 | Biosynthesis of amino acids | 3 of 66 | 1.21 | 0.53 | 0.0262 |
| dme01100 | Metabolic pathways | 10 of 1111 | 0.51 | 0.35 | 0.0262 |

**Biological Process**

| GO-term | Description | Count in Network | Strength | Signal | False Discovery Rate |
|---|---|---|---|---|---|
| GO:0006357 | Transcription initiation from RNA polymerase II promoter | 4 of 48 | 1.89 | 1.29 | 0.0016 |
| GO:0030490 | Maturation of SSU-rRNA | 3 of 49 | 1.76 | 0.91 | 0.0109 |
| GO:0016070 | RNA metabolic process | 8 of 861 | 0.94 | 0.73 | 0.002 |
| GO:0090304 | Nucleic acid metabolic process | 2 of 1143 | 0.86 | 0.68 | 0.002 |
| GO:0010467 | Gene expression | 8 of 1138 | 0.82 | 0.58 | 0.008 |
| GO:0043170 | Macromolecule metabolic process | 11 of 3264 | 0.5 | 0.34 | 0.0253 |
| GO:0044738 | Primary metabolic process | 17 of 4113 | 0.43 | 0.3 | 0.0007 |

**Cellular Component**

| GO-term | Description | Count in Network | Strength | Signal | False Discovery Rate |
|---|---|---|---|---|---|
| GO:0016591 | RNA polymerase II, holoenzyme | 4 of 71 | 1.72 | 1.45 | 0.00041 |
| GO:0090575 | RNA polymerase II transcription regulator complex | 4 of 111 | 1.53 | 1.27 | 0.00068 |
| GO:0030684 | Preribosome | 3 of 69 | 1.61 | 1.06 | 0.0037 |
| GO:1990234 | Transferase complex | 6 of 491 | 1.06 | 0.89 | 0.00084 |
| GO:0005669 | Transcription factor TFIID complex | 2 of 26 | 1.85 | 0.8 | 0.0208 |

**Figure S2. PPI Network of the 271 *Drosophila* Genes**

A) using all datasets with a confidence score of 0.4. Nodes represent individual proteins, and edges represent predicted functional associations. B) Gene Ontology (GO) terms identified in the enrichment analysis.

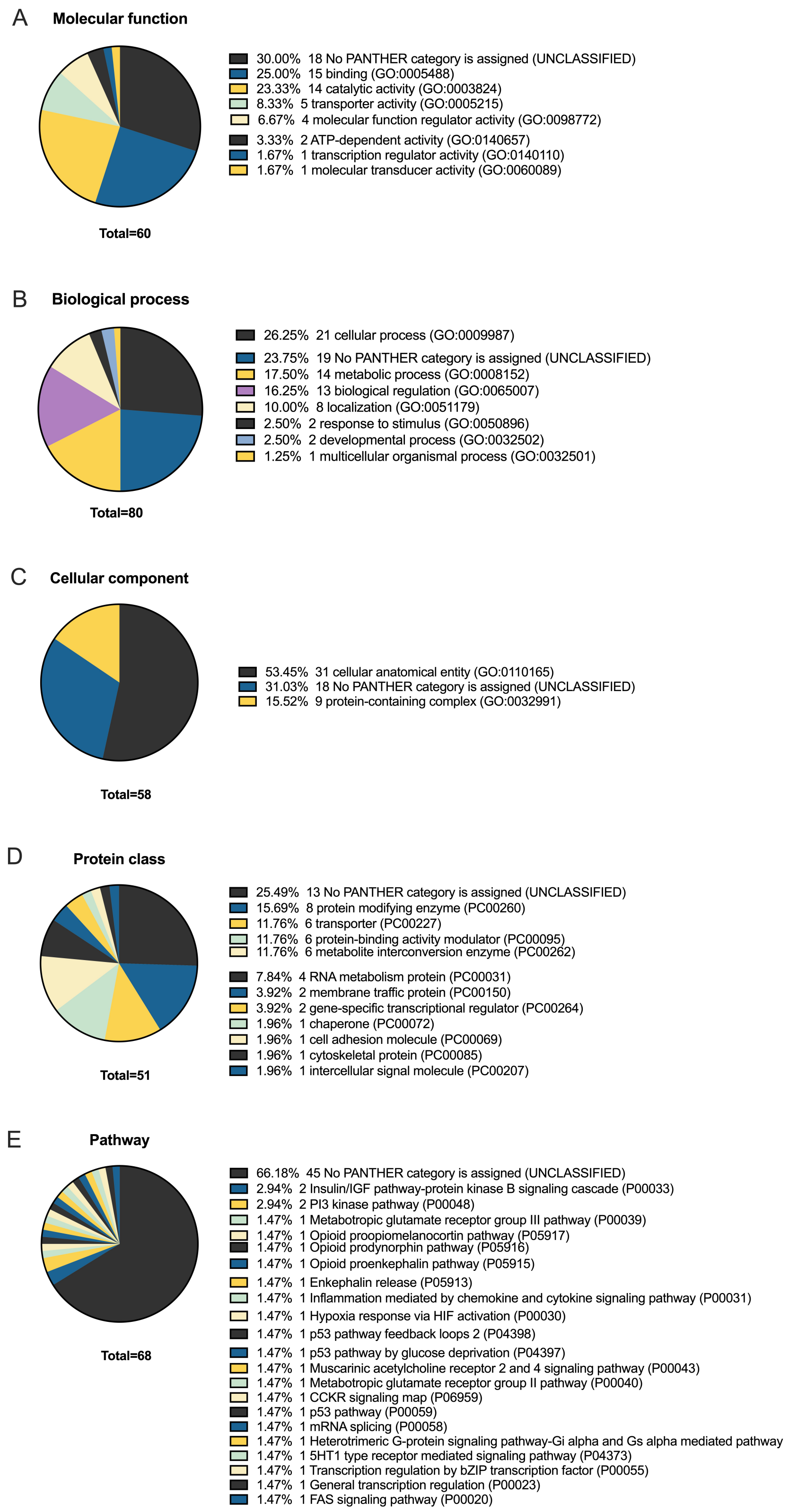
A
Molecular function
30.00% 18 No PANTHER category is assigned (UNCLASSIFIED)
25.00% 15 binding (GO:0005488)
23.33% 14 catalytic activity (GO:0003824)
8.33% 5 transporter activity (GO:0005215)
6.67% 4 molecular function regulator activity (GO:0098772)
3.33% 2 ATP-dependent activity (GO:0140657)
1.67% 1 transcription regulator activity (GO:0140110)
1.67% 1 molecular transducer activity (GO:0060089)
Total=60
B
Biological process
26.25% 21 cellular process (GO:0009987)
23.75% 19 No PANTHER category is assigned (UNCLASSIFIED)
17.50% 14 metabolic process (GO:0008152)
16.25% 13 biological regulation (GO:0065007)
10.00% 8 localization (GO:0051179)
2.50% 2 response to stimulus (GO:0050896)
2.50% 2 developmental process (GO:0032502)
1.25% 1 multicellular organismal process (GO:0032501)
Total=80
C
Cellular component
53.45% 31 cellular anatomical entity (GO:0110165)
31.03% 18 No PANTHER category is assigned (UNCLASSIFIED)
15.52% 9 protein-containing complex (GO:0032991)
Total=58
D
Protein class
25.49% 13 No PANTHER category is assigned (UNCLASSIFIED)
15.69% 8 protein modifying enzyme (PC00260)
11.76% 6 transporter (PC00227)
11.76% 6 protein-binding activity modulator (PC00095)
11.76% 6 metabolite interconversion enzyme (PC00262)
7.84% 4 RNA metabolism protein (PC00031)
3.92% 2 membrane traffic protein (PC00150)
3.92% 2 gene-specific transcriptional regulator (PC00264)
1.96% 1 chaperone (PC00072)
1.96% 1 cell adhesion molecule (PC00069)
1.96% 1 cytoskeletal protein (PC00085)
1.96% 1 intercellular signal molecule (PC00207)
Total=51
E
Pathway
66.18% 45 No PANTHER category is assigned (UNCLASSIFIED)
2.94% 2 Insulin/IGF pathway-protein kinase B signaling cascade (P00033)
2.94% 2 PI3 kinase pathway (P00048)
1.47% 1 Metabotropic glutamate receptor group III pathway (P00039)
1.47% 1 Opioid proopiomelanocortin pathway (P05917)
1.47% 1 Opioid prodynorphin pathway (P05916)
1.47% 1 Opioid proenkephalin pathway (P05915)
1.47% 1 Enkephalin release (P05913)
1.47% 1 Inflammation mediated by chemokine and cytokine signaling pathway (P00031)
1.47% 1 Hypoxia response via HIF activation (P00030)
1.47% 1 p53 pathway feedback loops 2 (P04398)
1.47% 1 p53 pathway by glucose deprivation (P04397)
1.47% 1 Muscarinic acetylcholine receptor 2 and 4 signaling pathway (P00043)
1.47% 1 Metabotropic glutamate receptor group II pathway (P00040)
1.47% 1 CCKR signaling map (P06959)
1.47% 1 p53 pathway (P00059)
1.47% 1 mRNA splicing (P00058)
1.47% 1 Heterotrimeric G-protein signaling pathway-Gi alpha and Gs alpha mediated pathway
1.47% 1 5HT1 type receptor mediated signaling pathway (P04373)
1.47% 1 Transcription regulation by bZIP transcription factor (P00055)
1.47% 1 General transcription regulation (P00023)
1.47% 1 FAS signaling pathway (P00020)
Total=68

**Figure S3. GO Term Analysis using PANTHER Classification system.**

The 271 Drosophila genes were categorized based on A) molecular function, B) biological process, C) cellular component, D) protein class, and e) pathways.
