## Supplementary figures and images for "TSC2 is a stress granule suppressor"

### Source Data for Western Blots

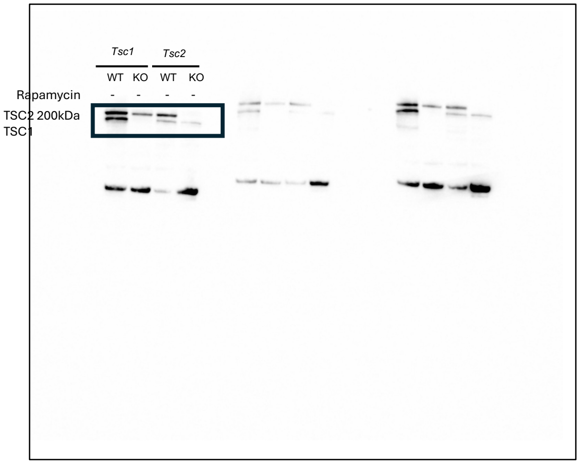

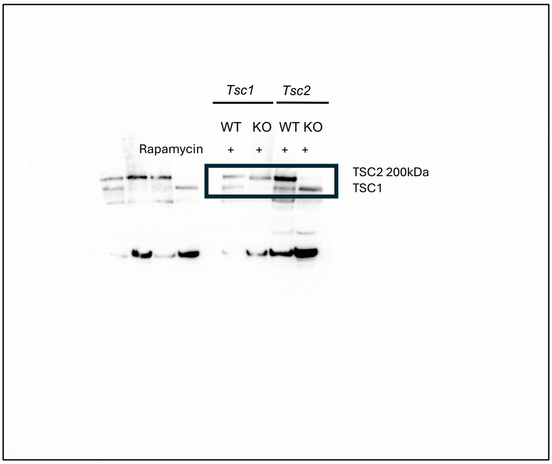

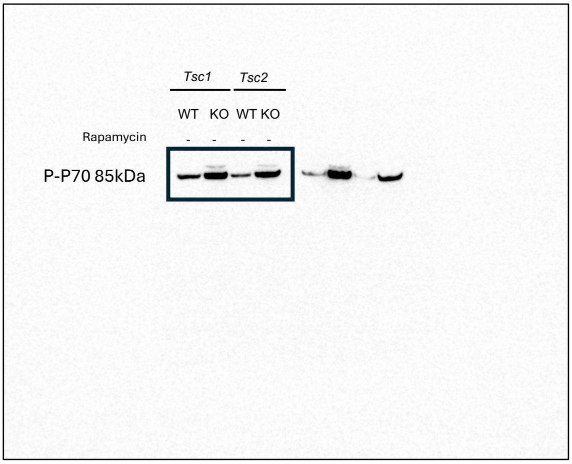

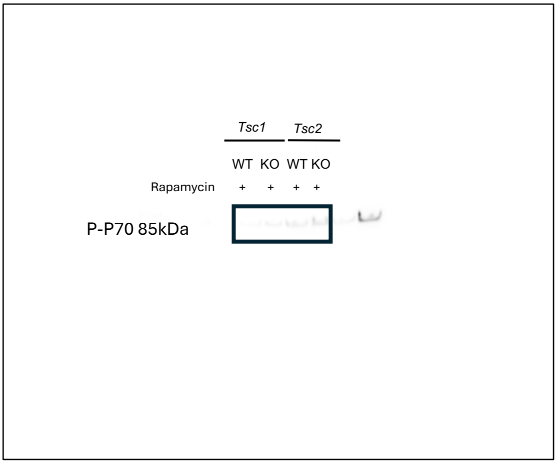

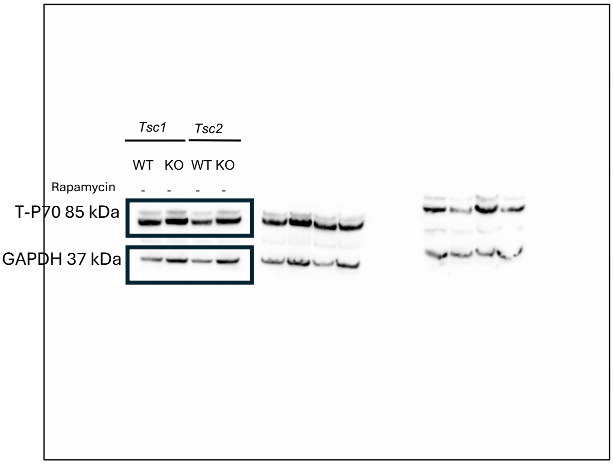

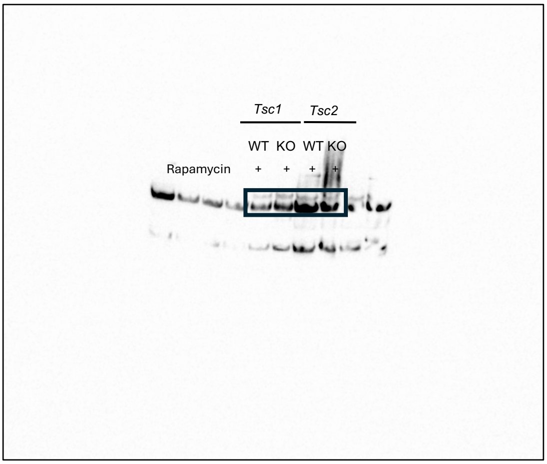

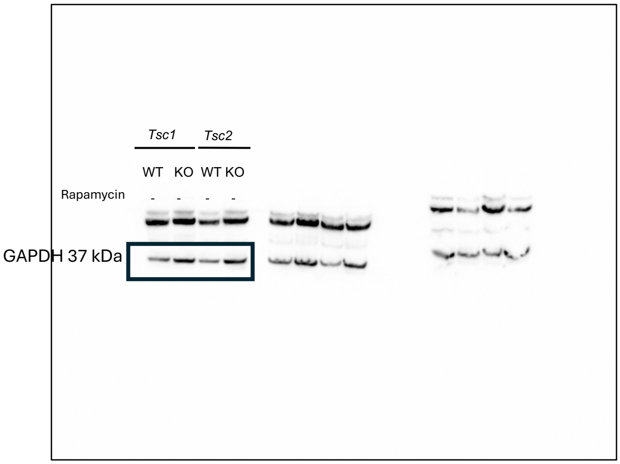

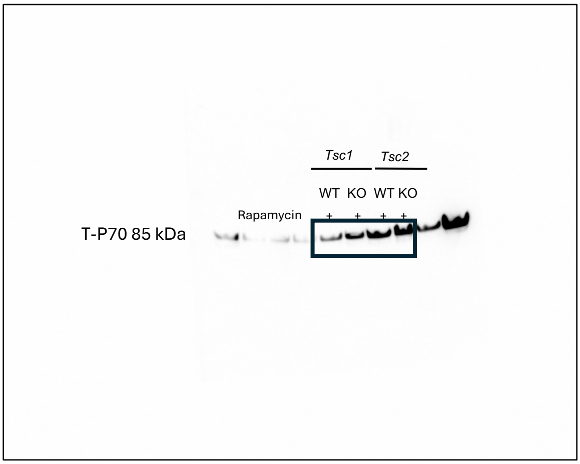

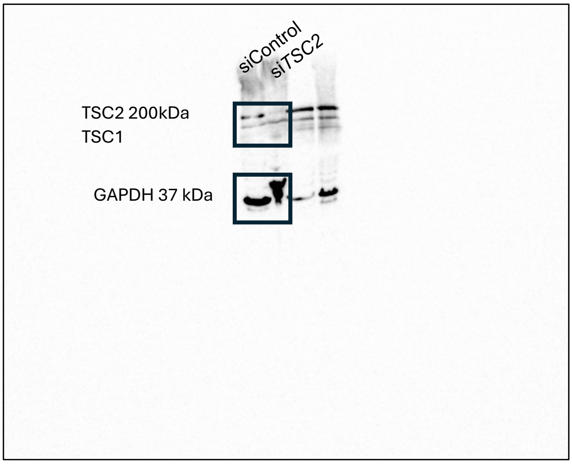

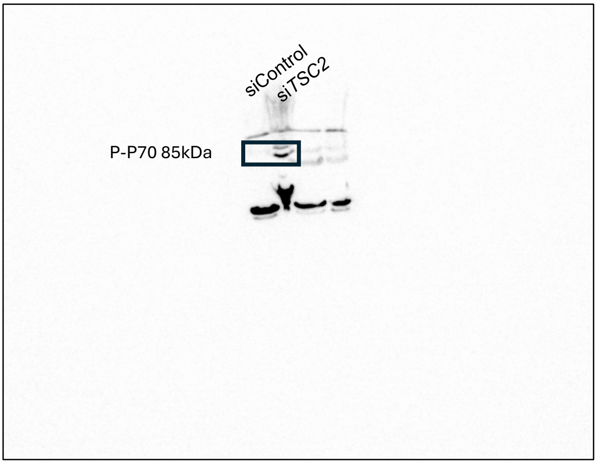

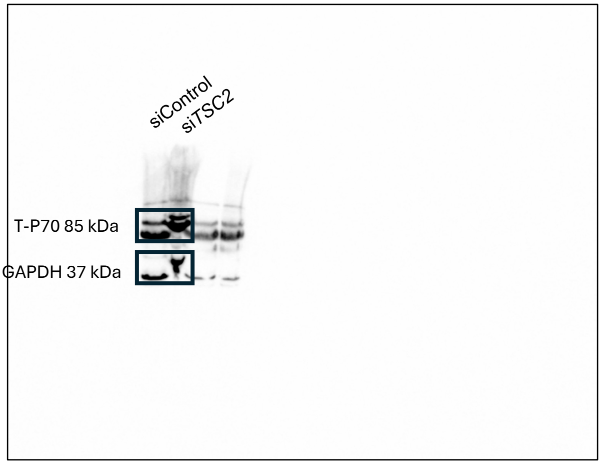

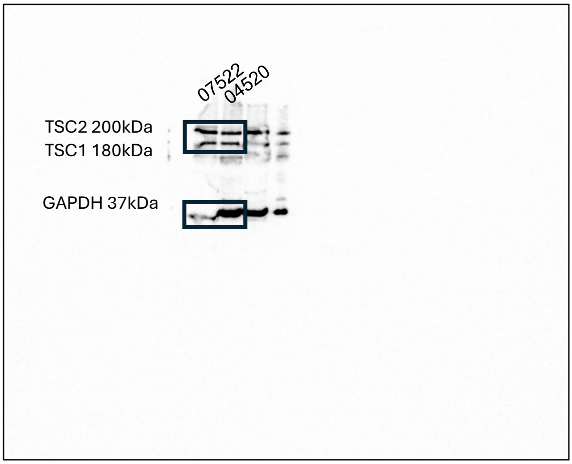

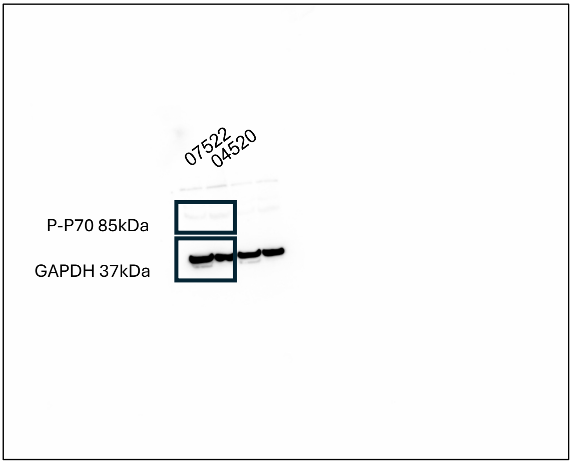

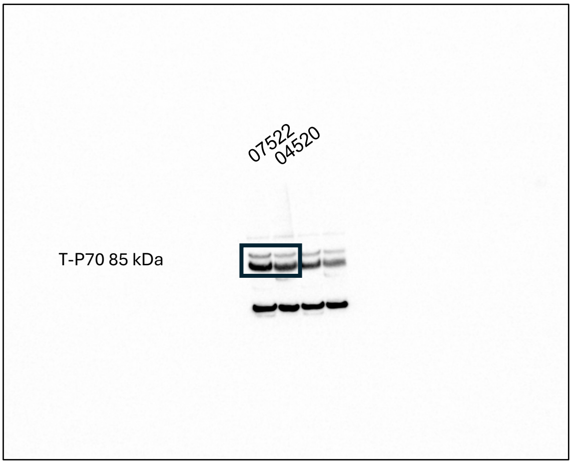
